# Refining mechanistic models to better predict larval and nymphal activity patterns of *Ixodes scapularis*

**DOI:** 10.64898/2026.08.26.747079

**Authors:** Stacy Mowry, T. Alex Perkins

**Affiliations:** University of Notre Dame

**Keywords:** ticks, life-cycle, phenology, Lyme disease, ordinary differential equations, nonsystemic

## Abstract

The black-legged tick (*Ixodes scapularis*), a key vector of Lyme disease, anaplasmosis, and babesiosis, exhibits regionally distinct patterns of seasonal activity driven by climate. Consequently, the relative timing of larval and nymphal activity varies across geographic locations, influencing pathogen transmission dynamics. Early-emerging nymphs may increase pathogen transmission, whereas early-emerging larvae may reduce transmission. In addition, synchrony between the two life-stages facilitates co-feeding transmission, which contributes to pathogen maintenance and coinfection risk. Temperature is thought to be an important driver of tick phenology, but existing mechanistic models that incorporate temperature fail to accurately capture the timing of larval and nymphal tick activity. To address this limitation, we developed a mechanistic model that includes two additional factors: humidity-dependent questing and low rates of overwinter development. To assess the value of these factors for explaining real-world patterns, we fitted alternative models to tick collection data from the National Ecological Observatory Network. In doing so, we found that explicitly incorporating humidity is necessary to reproduce observed tick phenology, with larval ticks being especially sensitive to relative humidity compared to other life stages. In addition, we found that accounting for humidity had a larger effect at Mid-Atlantic sites than at Northeastern sites, underscoring the importance of region-specific interactions between temperature and humidity in shaping *I. scapularis* phenology. By more accurately capturing tick seasonality compared to existing mechanistic models, our model illustrates the importance of accounting for factors beyond temperature for investigating how climate variability influences seasonal tick activity and pathogen transmission.

## 2 Introduction

The black-legged tick (*Ixodes scapularis*) is the primary vector for tick-borne diseases in the United States (Wolf et al., 2020), transmitting Lyme disease, anaplasmosis, ehrlichiosis, and babesiosis. Its geographic range extends from the eastern coast to eastern North Dakota and Texas (Eisen & Eisen, 2023), and its activity patterns vary both spatially and temporally. These differences in seasonal activity largely arise in response to climatic variation (Qviller et al., 2014). Understanding the relationship between climate patterns and tick activity is essential to accurately predict seasons of increased tick-borne disease risk within the context of a changing climate.

Seasonal activity of *I. scapularis* depends on the timing of tick development and questing. The tick passes through larval, nymphal, and adult stages, each requiring a blood meal before molting. The rate and timing of interstadial development is driven by both temperature-dependent developmental quiescence, with warmer temperatures generally accelerating development nonlinearly (Branagan, 1973; Chilton & Bull 1994, Peavey & Lane, 1996; Randolph, 1997; Randolph et al., 2002, Ogden et al., 2004), and temperature-independent diapause. Diapause is a genetically determined trait that temporarily halts development or activity in response to decreased day length (Belozerov et al., 2002), helping synchronize the tick life-cycle with generally favorable seasons (Gray et al., 2016).

The relative activity periods of larval and nymphal ticks can critically influence the transmission dynamics of tick-borne pathogens, and thus human risk of tick-borne disease. For example, the enzootic transmission cycle of *Borrelia burgdorferi*, which causes Lyme disease, depends on transmission between reservoir hosts and larval and nymphal *I. scapularis*. Transmission risk rises when nymphs feed earlier than larvae, since infected nymphs can infect hosts that later transmit the pathogen to larvae (Yuval & Spielman, 1990; Kurtenbach et al., 2006; Goethert et al., 2023). Because some host infections clear quickly, or wane over time (Donahue et al., 1987), a shorter gap between nymphal and larval feeding increases transmission. In addition to the sequential transmission pathway, synchrony in larval and nymphal activity, which is also influenced by climatic variation (Levi et al., 2015; Gatewood et al., 2009; Sambado et al., 2024), offers another mechanism for pathogen transmission. When both larval and nymphal ticks feed simultaneously on the same host, non-systemic transmission, which is when a pathogen spreads directly from an infected tick to an uninfected tick without amplification by the host (Randolph and Nuttall, 1996), can occur. Feeding synchrony can sustain transmission (Voordouw, 2015), particularly for rapidly cleared Borrelia strains (States et al., 2017). Synchrony may also impact the prevalence of coinfection in ticks and thus coinfection in humans, which can negatively impact human health by decreasing treatment efficacy (Diuk-Wasser et al., 2016) or increasing symptom severity (Thomas et al., 2001) or duration (National Institute of Allergy and Infectious Diseases). Together, these observations highlight the importance of accurately predicting the seasonal timing of larval and nymphal activity, the degree of synchrony between stages, and understanding the influence of climate on tick phenology.

One way to understand tick life cycles is through mechanistic modeling with ordinary differential equation (ODE) models. These models offer a tractable framework for predicting seasonal tick activity, quantifying disease risk across space and time, and assessing the effects of interventions and environmental change. When fitted to data, they can also recover biological rates and processes that are difficult to measure directly. ODEs have been widely used to examine how temperature shapes developmental thresholds and the basic reproduction number (*R*_0_) for tick (Wu et al., 2010) and pathogen establishment (Lou et al., 2014), and to analyze sensitivity to host population size (Lou et al., 2014), climate (Wallace et al, 2019; Husar et al., 2024, Winter et al., 2020; Wu et al., 2013), and land cover (Winter et al., 2020).

Although widely used, existing *I. scapularis* life-cycle models consistently fail to reproduce observed seasonal activity patterns without modification (Winter et al., 2020; Wu et al., 2013). One common workaround is to impose phenomenological seasonal forcing that restricts activity to fixed periods of the year (Winter et al., 2020; Husar et al., 2024). Although this can replicate local patterns, it decouples tick activity from environmental variation, limiting the ability to capture interannual variability, anomalous seasons, and responses to climate change. Alternatively, some models accumulate daily temperature-dependent development rates until a threshold is reached, where the development rate is defined as the reciprocal of the number of days required to reach that threshold (Wu et al., 2013; Lou et al., 2014). Even though this cumulative approach allows development to vary with temperature, it is inherently problematic in that future temperatures inform development rates before those temperatures are actually experienced by the organism. It also results in development rates being averaged across periods of weeks or months, which reduces the ability of models to capture nuances of seasonal timing.

In this study, we address limitations of existing mechanistic models by incorporating temperatures into tick life-history parameters instantaneously and by incorporating two additional hypothesized drivers of tick phenology into an existing ODE tick life-cycle framework (Ogden et al., 2005). The first mechanism, humidity-dependent questing, is well supported by empirical evidence that questing activity is constrained by desiccation risk (Fryxell et al., 2015; Stafford, 1994; Ginsberg et al., 2017). The second mechanism, a minimum larval-to-nymphal overwinter development rate, is based on the hypothesis that the buffered microclimates beneath snow or leaf litter (Volk et al., 2022; Linske & Williams, 2019) allow for development to proceed at a low level throughout the winter. This contrasts with models that completely pause development during winter months.

To assess the value of these factors for explaining real-world patterns of tick phenology, we incorporated both into an existing ODE tick life-cycle model and fitted it to seasonal tick drag data from the National Ecological Observatory Network (NEON), which provides standardized, longterm tick and environmental data from a number of sites across the United States. We then used model comparison to evaluate whether inclusion of these mechanisms improved model predictions of the seasonal timing and synchrony of larval and nymphal activity. Our overarching goal was to develop a climate-driven tick life-cycle model that improves predictive accuracy of tick seasonality.

## 3 Materials and methods

### 3.1 Overview

We modified a mechanistic, stage-structured model of the *Ixodes scapularis* life cycle to incorporate two hypothesized drivers of phenology: humidity-dependent questing behavior and a minimum larval-to-nymphal overwinter development rate. The model was formulated to allow parameter estimates to support the null effect of each hypothesized mechanism. We first fitted the model to larval and nymphal activity data from three NEON sites to estimate parameters governing these processes. To isolate the influence of mechanistic parameters from the influence of the prior on mechanistic parameter estimation, we performed a sensitivity analysis by visualizing how individual parameter perturbations affected the model likelihood. We then evaluated the performance of three model variants: a baseline “no-effect” model, a “prior-effect” model, and a “posterior-effect” model. Finally, we assessed each model’s ability to predict daily activity and seasonal synchrony within the NEON dataset.

### 3.2 Model description

Our model describes the life-cycle of *I. scapularis*, based on the model introduced by Ogden et al. (2005) (Figure 1). The original model consists of twelve discrete tick life stages, and ticks transition through these life-stages based on temperature-dependent development rates *ρ*(*T* (*t*)), host attachment rates *λ* (*H*) (which are modulated by host density, *H*), and temperature-dependent questing probabilities *θ* (*T* (*t*)). This original model also incorporates seasonal diapause behavior driven by changing day length and climate, where the temperature-dependent development rates *ρ*(*T* (*t*)), are set to 0 after the summer solstice (*t >* 180) or if ambient conditions fall below freezing (*T* (*t*) *≤* 0). Further detail on the full model is provided in the Supplement and all parameters are described in Table S1 and Table S2. Here, we focus on the modifications we made to the original model and their justifications.

**Figure 1.**
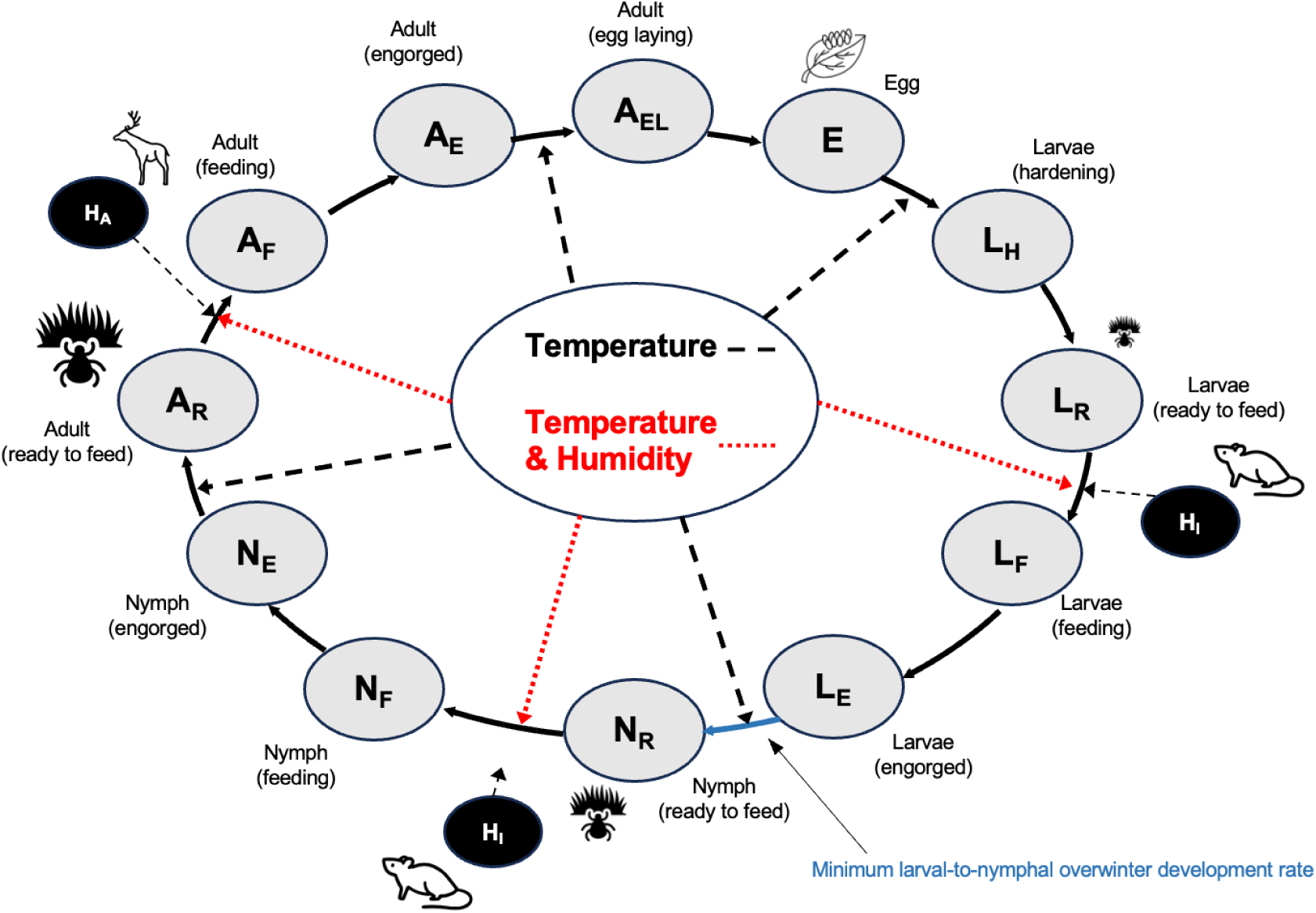
Graphical depiction of tick life-cycle model. Our first modification: humidity dependent questing is indicated in red. Our second modification: minimum larval-to-nymphal overwinter development rate is indicated in blue.

### 3.3 Model modifications

To evaluate whether additional biological processes could explain observed variation in seasonal tick activity, we introduced two hypothesized mechanisms: humidity-dependent questing and a minimum larval-to-nymphal overwinter development rate, into the original model. We estimated parameters associated with each hypothesized mechanism simultaneously to prevent parameter confounding. Although a sequential single-mechanism approach may produce a more parsimonious fit, our objective was to isolate the relative contributions of the individual biological mechanisms.

Questing success is strongly influenced by relative humidity, with desiccation risk limiting activity above the leaf litter (Stafford, 1994; Fryxell et al., 2015). Based on laboratory studies, nymphal ticks remain viable at relative humidities above 80%-85% (Fryxell et al., 2015), whereas lower relative humidity increases mortality and reduces questing success. Larvae are even more sensitive (Fryxell et al., 2015), requiring higher relative humidity for viability. To account for these effects, we modified the baseline model such that the probability of tick questing is shaped by relative humidity *M*(*t*) according to a sigmoid function,

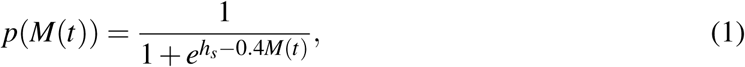

where *h_s_*determines the midpoint of the curve for each life stage *s*. We modeled the larval and adult midpoints as functions of the nymphal midpoint as *h^’^_l_ = h_n_ + h_l_* and *h^’^_a_ = h_n_ + h_a_* respectively. This allowed us to remove correlation between estimate *h_l_* and *h_a_*, removing any correlation between the effect of humidity on various lifestages, and improving MCMC sampling. For simplicity, we use *h_l_* and *h_a_* to represent *h^’^_l_* and *h^’^_a_* in our full model notation. Humidity did not directly affect mortality, consistent with ticks taking refuge beneath leaf litter. However, humidity indirectly affects mortality by maintaining ticks in a questing state for a longer period of time, which has a higher daily mortality rate than engorged stages. All thresholds were estimated by fitting model predictions to seasonal tick drag counts. We then redefined the questing function in our model from *θ* (*T* (*t*)) to *θ* (*T* (*t*)*, p*(*M*(*t*))). All model parameter values are provided in Table S1 and Table S2.

Previous modeling studies have documented a consistent delay in the predicted onset of nymphal questing when using laboratory-derived, temperature-development relationships (Winter et al., 2020; Wu et al., 2013). We hypothesized that such a delay may be a result of models underestimating development that occurs at air-temperatures below 0 °C. Mechanistically, we hypothesized that overwintering ticks may experience microclimates beneath snowpack or leaf litter, or occasional warm temperatures, that permit a small but nonzero amount of development even when daily average air temperature is below the developmental threshold. Given the relatively long duration of winter, these small rates could potentially add up to be impactful on tick phenology in the spring. We therefore introduced a modified larval-to-nymphal development function into our model, *f_ln_*(*T* (*t*)), that imposes a minimum development rate such that

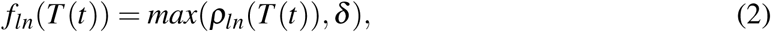

where *ρ_ln_*(*T* (*t*)) is the temperature-dependent larval-to-nymphal development rate as defined by Ogden et al., (2005) and *δ* is the estimated minimum larval-to-nymphal development rate. To improve algorithmic convergence we reparameterized 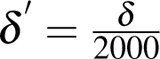 (again we use *δ* to represent *δ ^′^* in our notation). The parameter *δ* was estimated via fitting model predictions to seasonal tick counts.

### 3.4 NEON Data

We selected National Ecological Observatory Network (NEON) sites within the Northeastern and Mid-Atlantic portions of the *I. scapularis* range that had at least three years of data with both larvae and nymphs present. Sites in the southern range were excluded because these populations exhibit distinct behavioral and genetic characteristics (Arsnoe et al., 2019; Sakamoto et al., 2014) that affect tick activity. This process yielded three sites: Harvard Forest (HARV) in the Northeast and the Smithsonian Conservation Biology Institute (SCBI) and Blandy Experimental Farm (BLAN) in the Mid-Atlantic. Our data spanned 2016 to 2023. To ensure the data accurately reflected the full seasonal cycle, we excluded years where sampling may have missed the beginning or end of a life stage’s activity period. We defined criteria for this exclusion based on proximity to the start and end of the drag dates. Specifically, if the tick count or density recorded within two weeks of the first or last sampling day exceeded 85% of the annual peak at that site, the site-year was removed. This prevented the inclusion of years where high activity levels at the start or end of the season suggested that the true activity peak may have occurred outside the sampling window.

### 3.5 Model fitting

For each life stage (*s*), NEON site (*η*), and year (*y*), we modeled the observed tick count data *C_sηty_* on day *t* as arising from a Dirichlet-Multinomial distribution such that

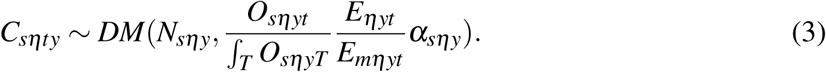

Here, *N_sηy_* is the total number of ticks of life stage *s* collected at site *η* over the year *y*. The expression 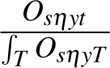 describes the model-predicted percent of tick activity at time *t* for ticks of life-stage *s* at site *η* in year *y*. The predicted percent activity was then scaled based on sampling effort on day *t* for each year and life-stage relative to the maximum sampling effort over all days of the sampling period yielding the full mean, 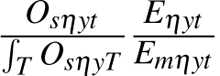. Finally, *α_sηy_* is the concentration parameter, which controls the concentration of the probability density around the mean. Our goal was to estimate the mechanistic parameters *h_s_* and *δ*, the humidity threshold for each life stage and larval to nymphal overwinter development rate, respectively. We also estimated the concentration parameters for each life-stage and site-year.

### 3.6 Bayesian inference

The ODE model was implemented in *odin2* (FitzJohn, 2025) and the R package *BayesianTools* (Hartig et al., 2023) was used for model fitting. Mechanistic parameters *h_sη_* and *δ_η_* were pooled across sites. The median prior on *h_sη_* was set to reflect reasonable certainty in a critical relative-humidity of 90%, which is consistent with existing system knowledge (Vail & Smith, 2002; Ostfeld & Brunner, 2015) (Figure2, Table S3). Priors on the relative humidity thresholds for larvae and adults were then formulated based on the expectation from system knowledge (Ginsberg et al., 2017) that larvae require a modestly higher relative humidity to remain active, whereas adults can quest at modestly lower relative humidities than immature stages (Figure 2; Table S3). Priors on the minimum larval-to-nymphal overwinter rate were formulated to reflect the expectation that development does not exceed the rates predicted by current laboratory-derived, temperature-driven development curves (Ogden et al., 2004) (Figure 3, Table S3). Concentration parameters were independent with lognormal (2,1) priors, reflecting our expectation of overdispersion in tick count data while constraining estimates to remain within realistic variability. Full prior specifications and prior predictive checks are provided in the Supplement. We ran three chains with 20,000 iterations and a burn-in period of 2,000 iterations. Model convergence was assessed using visual inspection of trace and density plots and verification that *R*^2^ values were *<* 1.1. Overall model fit, apart from our specific evaluation metrics, was assessed by calculating average *R*^2^ for each model over all site-years.

**Figure 2.**
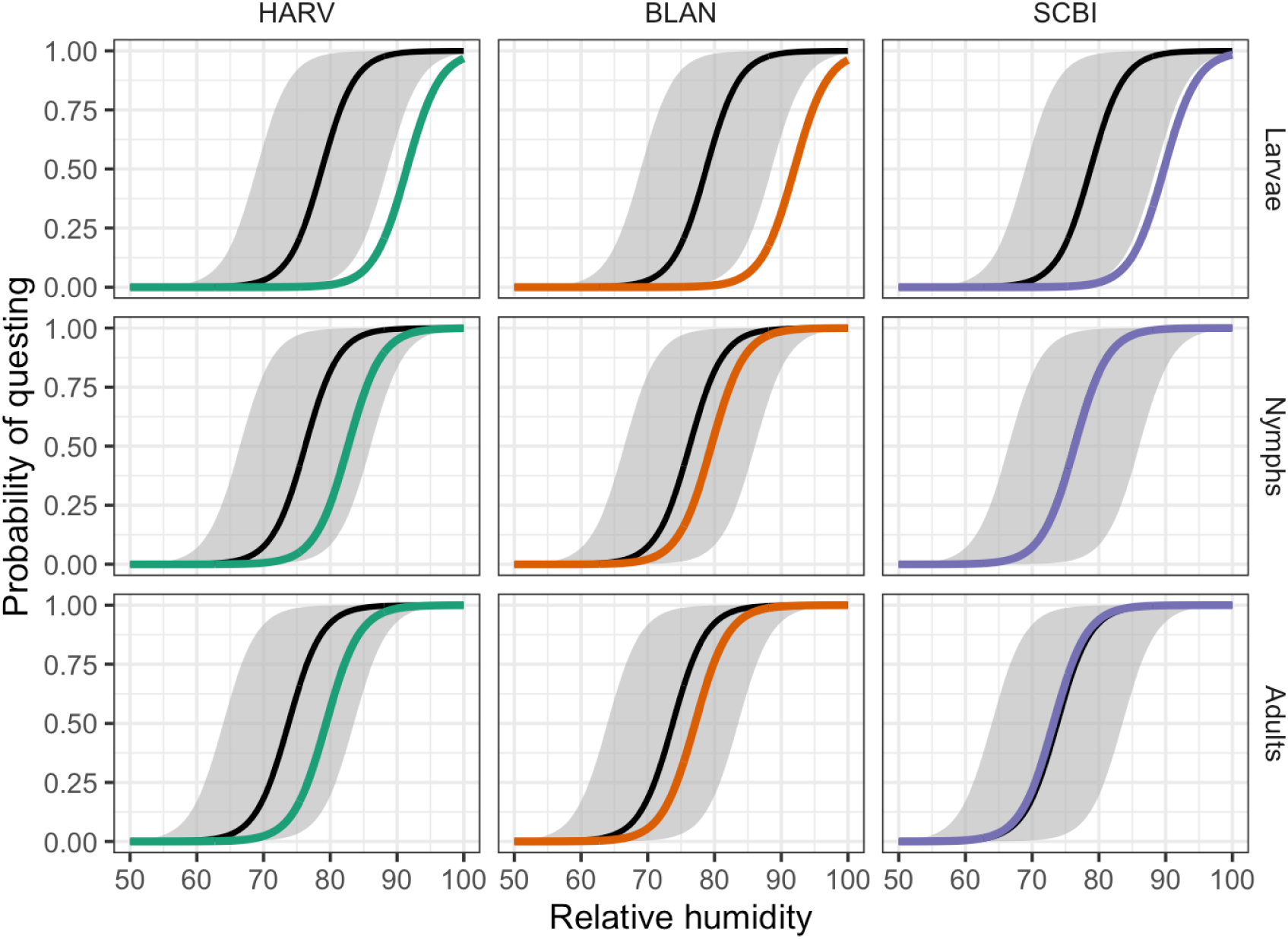
Estimates of the effect of relative humidity on the probability of questing across sites and life-stages. Thick black lines and grey shaded areas represent median prior expectations and 95% credible intervals (CIs), respectively. Thick colored lines indicate median posterior predictions. Results suggest larvae are more sensitive to desiccation than expected a priori, demonstrated by the median effect falling outside the prior predictive envelope across all sites.

**Figure 3.**
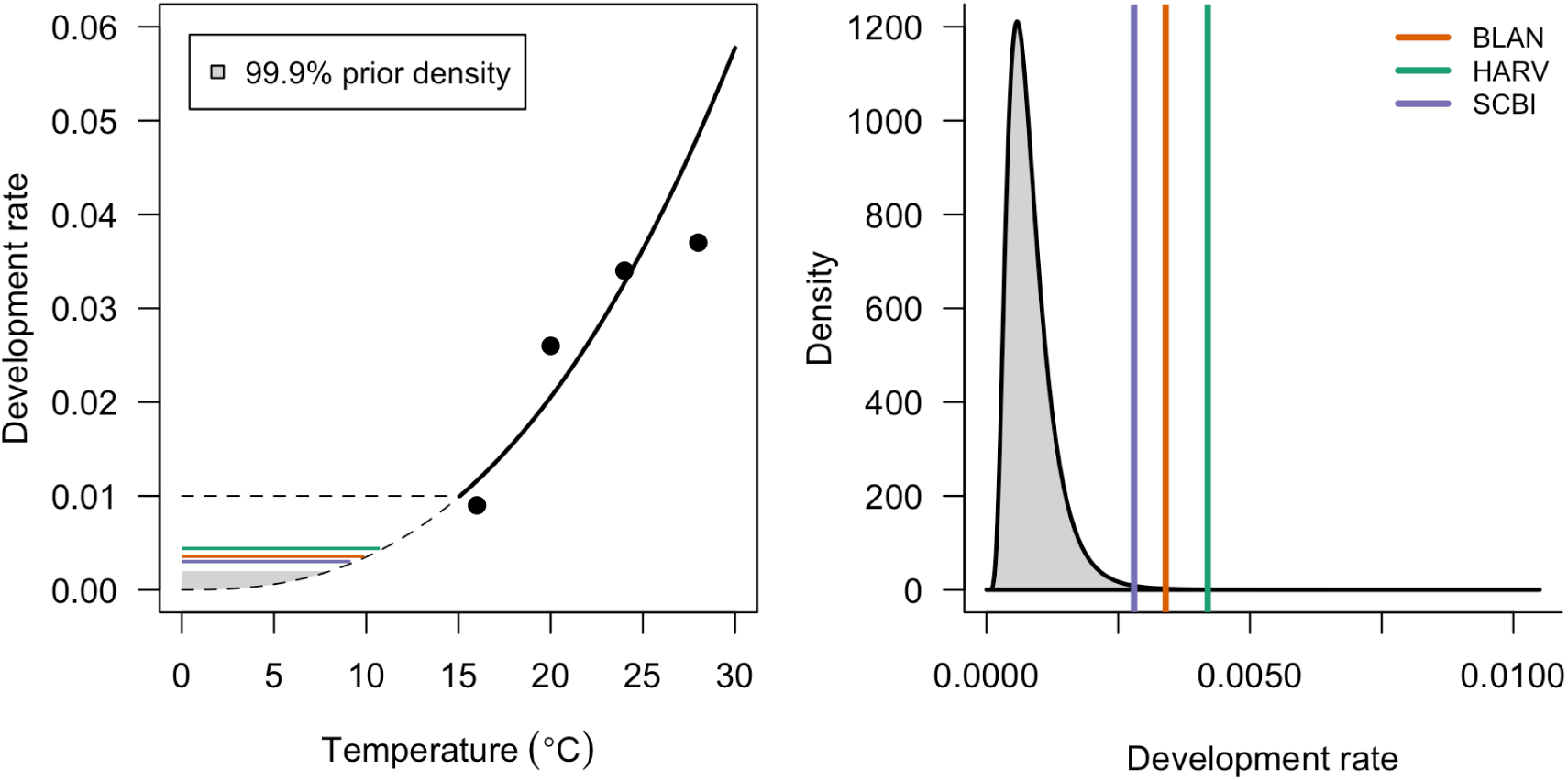
Posterior estimates of minimum overwinter larval-to-nymphal development across sites compared with the prior distribution. In the first panel, the solid curve shows the temperature-dependent larval-to-nymphal development rate estimated by Ogden et al., (2004) and the data which informed the relationship (dots). Dashed lines indicate the 99.99% prior credible interval for the estimated minimum larval-to-nymphal development rate. The second panel highlights the estimated posterior medians for each site relative to the prior distribution.

**Figure 4.**
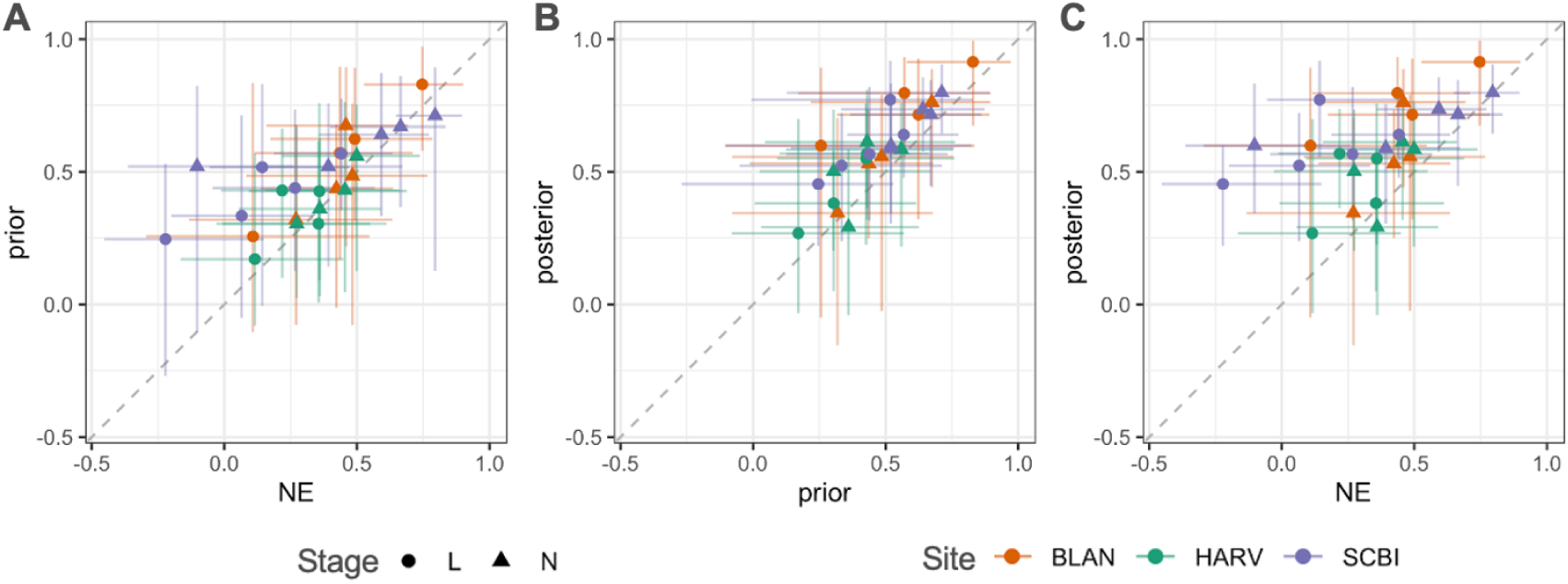
Comparison of model performance via Spearman rank correlations. Scatterplots represent site-year comparisons of rank correlations between empirical data and model-predicted larval and nymphal counts. Panels compare (A) no-effect (NE) vs. prior models, (B) prior vs. posterior models, and (C) NE vs. posterior models. The dashed diagonal represents the 1:1 line of equivalent performance. Error bars indicate 95% posterior predictive intervals for the rank correlation. The results show a stepwise improvement in predictive accuracy from the no-effect model to the prior, with the posterior model achieving the highest overall correlation.

### 3.7 Model comparison

To evaluate whether the inclusion of hypothesized mechanisms improved the model’s ability to reproduce observed tick phenology, we assessed model predictions using the seasonal timing of larval and nymphal activity and the degree of temporal synchrony between life stages. These metrics were selected because they correspond directly to the biological processes hypothesized to influence pathogen transmission dynamics and climate-driven variation in tick activity. We conducted validation against the NEON training data by comparing predicted seasonal timing and synchrony to observed activity patterns across three model variants: a no-effect model (excluding hypothesized mechanisms), a prior-effect model (fixing parameters at median prior expectations), and a posterior-effect model (using median posterior estimates after data fitting). The prior-effect model served as a test of the hypothesized mechanisms independent of data fitting. To assess overfitting, we calculated the DIC for each model, which assesses model fit while penalizing for model complexity.

#### 3.7.1 Isolating the effect of data on mechanistic parameter estimates

To distinguish between the influence of the data and the influence of the prior distributions on our posterior estimates, we performed a one-at-a-time sensitivity analysis for each mechanistic parameter. To do so, we calculated the log-likelihood across a range of parameter values while fixing other mechanistic parameters at their maximum *a posteriori* (MAP) estimate. Auxiliary parameters were fixed at their medians and one standard deviation above and below their medians, as determined by prior distributions. Alignment between the peak of the marginal likelihood profile and the full posterior indicates that the data supported the biological hypothesis represented by the prior. Conversely, divergence between the marginal likelihood peak and the posterior estimate suggests that the posterior is influenced by the prior distribution and that the data did not support the biological hypothesis. This sensitivity check ensured that our final parameter estimates, and our interpretation of them in light of our hypotheses, were driven by empirical observation rather than prior assumptions.

#### 3.7.2 Evaluating the seasonal activity of larvae and nymphs

To assess whether the model correctly reproduced the seasonal timing of larval and nymphal activity, we compared simulated phenology from the no-effect, prior-effect, and posterior-effect models to empirical phenology patterns derived from NEON tick-drag data at three sites. For each site, we generated environmental inputs by fitting periodic splines to annual temperature and humidity data. Missing values were imputed using the mean for that calendar day across available years. We then ran the model to equilibrium and simulated tick counts for each life stage for every site-year. For each model variant, model performance was quantified by calculating the Spearman correlation between normalized predicted tick feeding activity (*L_F_* for larvae and *N_F_* for nymphs) on each day that dragging was conducted and normalized drag counts from observed NEON tick-drag data. Higher correlation values indicate better phenological agreement between the model and empirical data. Spearman correlation was used because our goal was to assess if our models could capture phenological timing rather than absolute abundance.

#### 3.7.3 Evaluating larval and nymphal activity synchrony

To evaluate whether the model captured the degree of temporal synchrony between larval and nymphal activity, we compared synchrony predicted by the no-effect, prior-effect, and posterior-effect models to synchrony estimated from empirical NEON data. Synchrony is a key ecological parameter because concurrent feeding of larvae and nymphs on the same host can facilitate non-systemic transmission and alter infection dynamics.

For each site, year, and life stage, we generated empirical daily activity curves by fitting Weibull distributions to NEON tick collection data using the *MASS* package (Venables & Ripley, 2002). We used nonlinear least squares to fit these distributions to daily tick counts, weighting the counts by the total area sampled; this ensured that days with smaller sampling efforts had a smaller impact on the estimated activity distribution. The resulting curves were then normalized so that area under the curve summed to one and represented the underlying “true” seasonal activity for use in subsequent synchrony analyses. To account for potential error in the fitted distribution, we quantified the uncertainty associated with these curves by applying a Laplace approximation to the Weibull parameters (Figure S14).

Next, we used each prior and posterior draw from our modified models to generate daily predictions of larval and nymphal feeding activity. Then, we fitted generalized additive models (GAMs) to each draw using the *gam* function in the *mgcv* package (Wood, 2011). GAMs were used in place of our model predictions solely to smooth curves and facilitate integration necessary for our synchrony calculation. The GAM estimates were not substantially different from our model estimates (Figure S5). The smoothed curves normalized to sum to one, consistent with our empirical activity curves.

Synchrony between larval and nymphal activity was quantified as the proportional area of overlap between their respective normalized activity curves (Gatewood et al., 2009), which captures the fraction of total seasonal activity occurring simultaneously across both life stages. For both model-derived predictions and empirically-fitted Weibull distributions, the synchrony *S* is calculated as

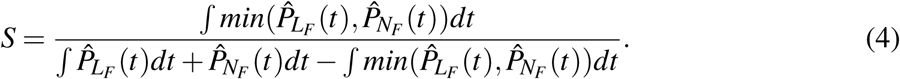

In this equation, P^_LF_ (*t*) and P^_NF_(*t*) represent the normalized larval and nymphal activity curves, respectively. For the mechanistic model predictions, *P* represents the smoothed GAMs; for the statistical models fitted to empirical data, *P* represents Weibull distributions.

## 4 Results

### 4.1 Model fit and convergence

All model parameters converged with Gelman-Rubin statistics *<* 1.1 and adequate mixing indicating chain convergence (Figure S4). The posterior model had an *R*^2^ of 0.304 (0.000618 – 0.838) compared to 0.239 (0.000281 – 0.770) for the prior model and 0.228 (0.000297 – 0.752) for the no-effect model. Evaluation of DIC revealed that the full model (DIC = 2,458) fit the data far better than the no-effect model (DIC = 2,660), even when accounting for additional model complexity, suggesting that the full model is not overfitted to the data.

### 4.2 Parameter estimation

#### 4.2.1 Humidity thresholds

We first estimated the effect of relative humidity on questing activity for each life stage. Effects were partially pooled across sites, reflecting a shared physiological response to humidity while allowing site-level differences associated with differences in regional climate. The relative humidity thresholds, defined as the relative humidity required for a 50% probability of questing, for larvae, nymphs, and adults, respectively, were: 91.5 (95% Bayesian Credibility Interval: 83.1 - 100), 82.7 (80.2 - 86.3), 79.4 (75.1 - 84.4) at HARV; 92.0 (82.1 - 100), 79.5 (75.4 - 84.1), and 77.1 (70.5 - 82.1) at BLAN; and 89.8 (83.8 - 100), 76.4 (74.6 - 78.0), and 73.2 (68.5 - 76.8) at SCBI (Figure 2, Figure S6). Based on comparison to the prior expectations, results indicated that larvae exhibited greater sensitivity to humidity than expected *a priori* at all sites, with more than 99% of larval posterior values being greater than the prior median for all sites. Estimates for the nymphal and adult thresholds were consistent with our prior expectations across sites.

#### 4.2.2 Minimum overwinter larval-to-nymphal development

As with the effect of relative humidity on questing, the minimum overwinter larval-to-nymphal development rate was partially pooled across sites, reflecting a shared physiological constraint on overwinter development while allowing site-level deviations associated with differences in regional climate and site-specific conditions. Across all sites, posterior median estimates were small but consistently shifted away from zero, exceeding the 95% prior credible interval at each site (HARV: 0.004 [0.0005 - 0.009], BLAN: 0.0034 [0.0002 - 0.008], SCBI:0.003 [0.0004 - 0.008]; Figure 3), indicating low but nonzero overwinter larval-to-nymphal development.

#### 4.2.3 Concentration parameters

Beyond estimating the parameters governing our biological mechanisms of interest, the model captured distinct aggregation patterns between life stages via the Dirichlet-multinomial concentration parameter. Larvae exhibited intense clustering (mean *α_l_* = 7.4), which corresponds to an intra-cluster correlation of approximately 0.119 and a variance inflation factor of 22 during peak collection events. Intra-cluster correlation is calculated as

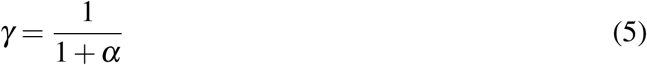

and variance inflation factor is calculated as

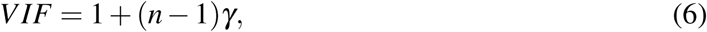

where *n* is the maximum larval or nymphal tick burden. A VIF of 22 indicates that larval variance was 22 times greater than expected under a standard multinomial distribution. Nymphs were distributed much more evenly (mean *α_n_*= 22.1), corresponding to a VIF of approximately 1.4 during peak collection. Full posterior distributions of concentration parameters across all specific site-years are available in Figure S7.

### 4.3 Model performance

#### 4.3.1 The effect of data on mechanistic parameters

Likelihood profile analysis revealed a strong data signal for the nymphal questing threshold (*h_n_*) across all sites and for the larval questing threshold (*h_l_*) at SCBI and BLAN. The weaker signal at Harvard Forest (HARV) likely reflects the site’s consistently high relative humidity during the larval questing period, which limits the environmental contrast needed to constrain this parameter. In contrast, the adult questing threshold was informed primarily by its prior distribution. This weak identifiability is structurally expected because the model was fitted exclusively to larval and nymphal field data. Adult *Ixodes scapularis* feed predominantly on deer (Spielman, 1985), which are poor reservoirs for *Borrelia burgdorferi*, and adult infection dynamics are therefore commonly omitted or simplified in transmission models (Ogden et al., 2013; MacDonald et al., 2021; Shauber & Ostfeld, 2002; Musa & Gumel, 2026; Haven & Park, 2012). Consequently, the limited data support for the adult questing threshold represents an acceptable limitation rather than a model deficiency. The close alignment between likelihood profile maxima and posterior parameter estimates further suggests strong agreement between the priors and the signal within available data (Figure S8).

#### 4.3.2 The seasonal activity of larvae and nymphs

To quantify model performance for predictions of seasonal tick activity, we calculated the rank correlation between normalized model predictions and normalized observed drag counts for each sampling day, averaging correlations across larval and nymphal life stages for each site-year. Across all site-years, mean rank correlation increased progressively from the no-effect model (0.35; 95% CI: *−*0.24 to 0.82) to the prior-effect model (0.46; 95% CI: *−*0.03 to 0.88) and was highest for the posterior-effect model (0.58; 95% CI: 0.10 to 0.92). These improvements are evident in the full activity curves, where the posterior model predicts distinct and separate peaks of tick activity for larvae and nymphs, consistent with the patterns observed in the data (Figure 5; Figure S9; Figure S10). The fact that the prior-effect model outperformed the no-effect model suggests that the inclusion of the hypothesized mechanisms correctly captures the fundamental drivers of phenology. The further improvement observed in the posterior-effect model indicates that the data allow for an empirical refinement of the magnitude of these effects in the field. This stepwise improvement confirms that the model’s predictive power is rooted in biological realism rather than statistical overfitting. Full prior and posterior distributions of activity curves, and uncertainty of the fitted Weibull distributions, are given in Figure S11-Figure S14. Rank correlations for each model, site, year, and life stage are presented in Tables S4, S5, and S6.

**Figure 5.**
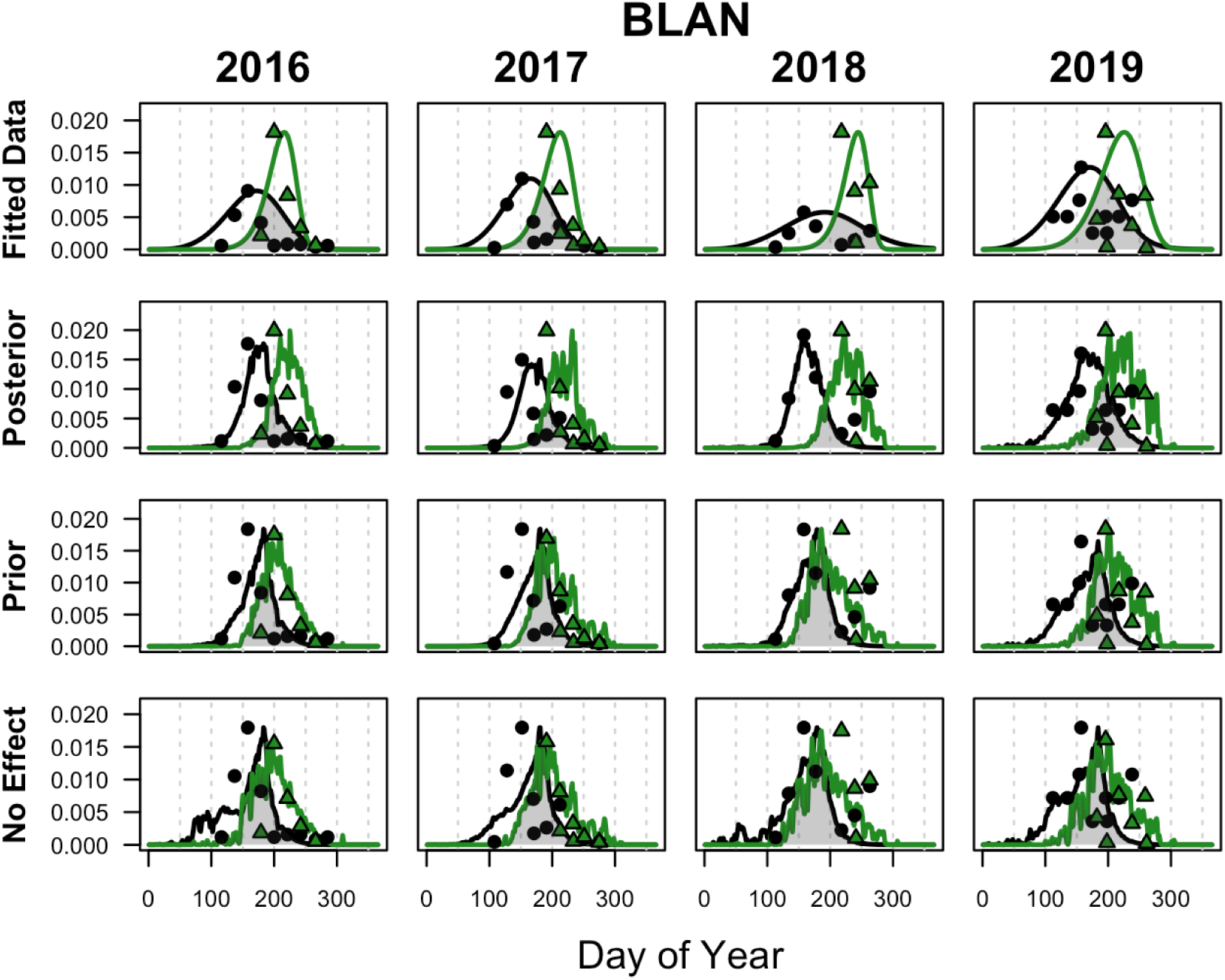
Comparison of data, statistical fit, and mechanistic model predictions for BLAN. In all panels, points represent observed field data and solid lines represent fitted activity curves for nymphs (black) and larvae (green). The top row displays empirical curves derived from statistical fits (Weibull distributions), representing the best estimate of true phenological peaks. Subsequent rows display median predictions from the posterior and prior models, alongside predictions from the no-effect model. The gray shaded regions emphasize seasonal synchrony.

#### 4.3.3 Larval and nymphal activity synchrony

Across site-years, incorporating the hypothesized mechanisms generally improved predictions of larval-nymphal synchrony relative to the no-effect model, with larger improvements seen at two of the three sites. At BLAN, 75% of prior draws and 69% of posterior draws were closer to the observed median synchrony than the no-effect model, while at SCBI the improvement was more pronounced (62% and 97%, respectively). In contrast, at HARV the proportion of draws closer to the observed median was lower (27% prior; 16% posterior). This pattern largely reflects that the no-effect model already produced accurate synchrony estimates for two of the three years (2018, 2021) at this site. The prior and posterior predictions in those years remained consistent with observations, as shown by overlap between the empirical and prior/posterior distributions (Figure 6). In the remaining HARV year (2017), the prior distribution improved model predictions compared to the no-effect model, but the posterior predictions diverged more strongly from the empirical estimate. This discrepancy, however, should be interpreted cautiously because our Weibull-derived empirical activity curves are influenced by the sparsity of NEON sampling days. Given the sensitivity of any single-year synchrony estimates to the empirically fitted activity distributions, overall patterns across sites and years provide a more robust indicator of model performance. Overall, prior and posterior synchrony distributions overlapped observed values, or improved estimates relative to the no-effect model, in all but one site-year, demonstrating clear improvement of synchrony predictions with the inclusion of hypothesized mechanisms. Compared to the prior model, the posterior model exhibited reduced variance compared to the prior, indicating increased precision of the biological mechanisms informed by the data (Table S7). Accurate synchrony predictions alone do not guarantee mechanistic validity, and only models that reproduce the underlying activity curves should be considered biologically supported. Importantly, the median activity curves demonstrate that the improved synchrony predictions from the posterior model arise from mechanistically realistic shifts in phenology (Figure 5).

**Figure 6.**
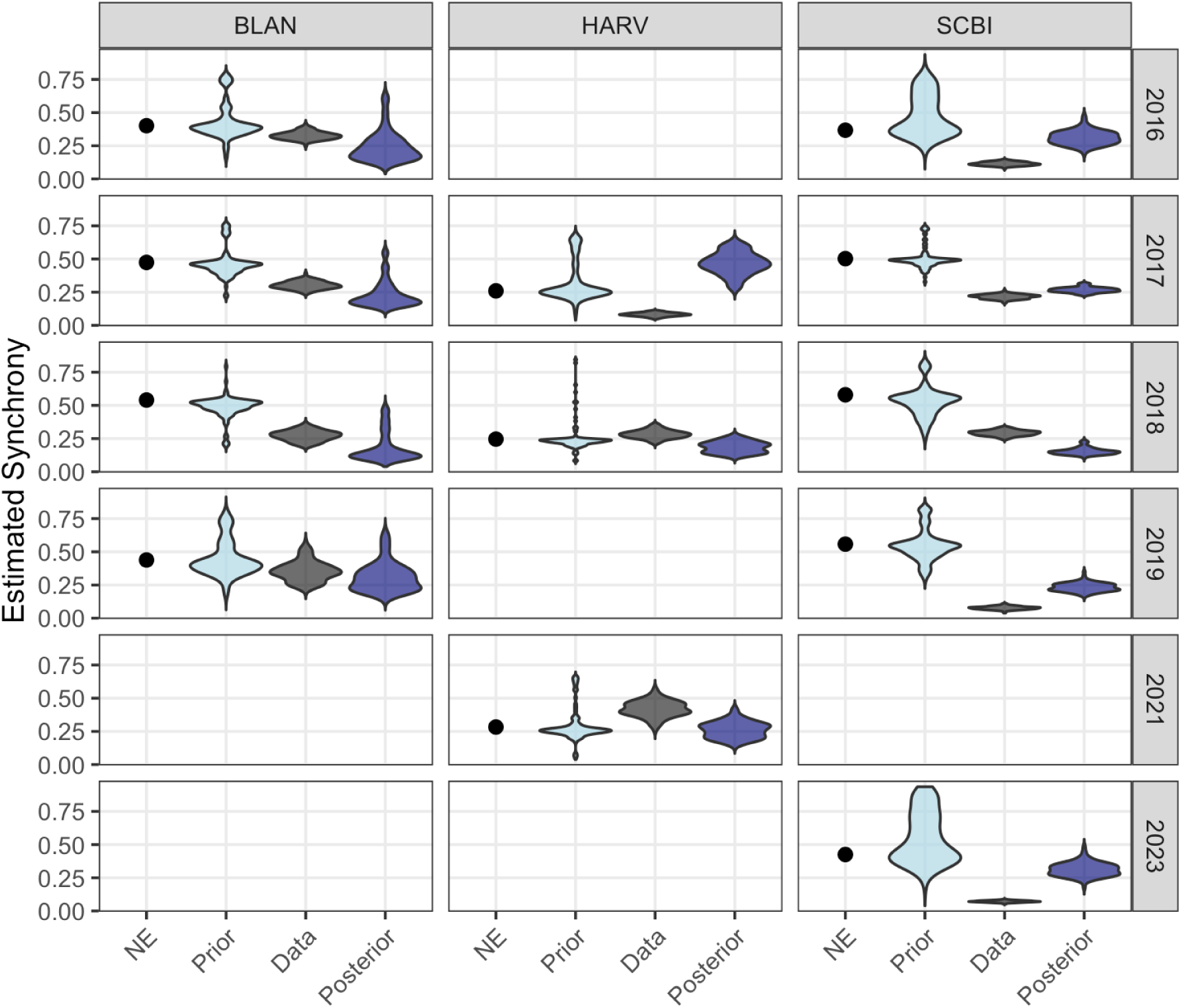
Distribution of synchrony predictions for each site year. Violin plots represent the predicted distributions of feeding synchrony for the posterior model (dark blue violin), prior model (light blue violin), no effect model (black dot) and empirical data (gray violin). Empirical synchrony was calculated by fitting Weibull distributions to larval and nymphal tick counts. For each model type, a greater degree of distributional overlap with the empirical data (gray) indicates better model prediction of seasonal synchrony.

#### 4.3.4 Effects of each parameter on model predictions

To isolate the contribution of each estimated mechanism, we generated model predictions using either the posterior estimate of the humidity effect alone or the posterior estimate of the minimum overwinter larval-to-nymphal development rate alone. Although the prior for minimum overwinter development concentrated most mass near zero, posterior estimates were consistently shifted away from zero across all sites, with posterior medians exceeding the 95% prior credible interval (Figure 3). Predictions from the development-only model closely resembled those of the no-effect model, whereas predictions from the humidity-only model closely matched those of the full posterior model. The primary difference introduced by the development-only model was a modest increase in spring nymphal activity relative to the no-effect model, with the largest difference at the coldest site, HARV (Figure 7). Across all sites, incorporating relative humidity narrowed the seasonal window of activity for both larvae and nymphs. This addition delayed larval peaks at all locations, with the most pronounced shifts occurring at BLAN and SCBI. The effect on nymphal phenology was site-specific: the nymphal peak was substantially accelerated at BLAN, modestly accelerated at SCBI, and unchanged at HARV.

**Figure 7.**
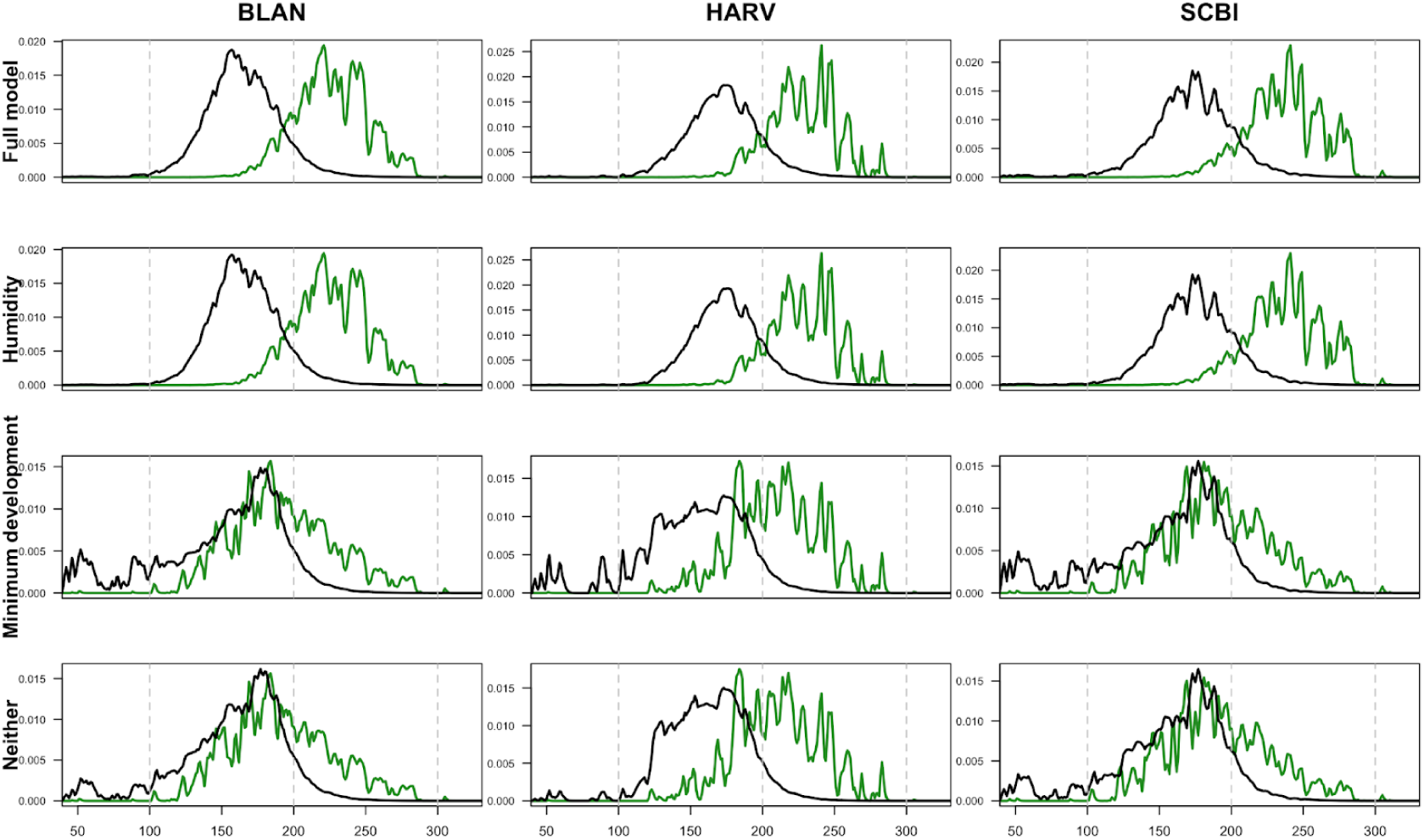
Influence of specific mechanisms on predicted seasonal phenology. Comparison of individual biological mechanisms and their impact on predicted larval (green) and nymphal (black) activity across all locations for 2018. Panels contrast the full posterior model (top row) and no-effect model (bottom row) against single-mechanism versions isolating humidity-dependent questing and minimum overwinter development. These comparisons demonstrate that shifts in the posterior and prior model behavior are primarily driven by the inclusion of humidity-dependent questing rather than inclusion of a minimum larval-to-nymphal development rate and that the effect of incorporating humidity is site-dependent, driven by the differences in each site’s unique climate profile.

## 5 Discussion

Our results demonstrate that incorporating two hypothesized biological mechanisms, informed directly from NEON data, improves the ability of a mechanistic model to reproduce larval and nymphal phenology relative to a temperature-only formulation. This improvement is supported by higher rank correlations between model predictions and observations and, in most cases, synchrony estimates that are closer to observed values. When we isolated the contribution of each mechanism, humidity-dependent questing emerged as the dominant driver of improved performance, whereas overwinter development, although consistently estimated as nonzero, produced only modest changes in seasonal activity.

A consideration when introducing and estimating additional parameters via model-fitting is the risk of overfitting. However, several lines of evidence demonstrate that our posterior estimates captures genuine information from the data. First, if the model were overfitted, we would expect the posterior formulation to improve predictions relative to the no-effect model, but the prior model, which incorporates the hypothesized mechanisms independently of the field data, to show no improvement. Instead, we demonstrated a clear, stepwise performance increase from the no-effect model, to the prior model, and finally to the posterior model, indicating that the biological mechanisms themselves, rather than arbitrary tuning to the data, are driving the improvements. Second, an overfitted model could yield parameter estimates that diverge from existing literature and lack ecological plausibility. In contrast, our posterior estimates for the nymphal humidity thresholds remained consistent with prior expectations and the divergent estimates for the larval threshold and the minimum overwinter development rate possess clear biological explanations. Finally, because the model was explicitly structured to allow the parameter estimates to support the null, “no-effect” model, we were able to isolate the data’s influence by calculating the likelihood of each parameter informed by the data alone. This technique demonstrated strong agreement with the full MCMC estimates, confirming that our parameters were driven by the empirical observations rather than based solely on our priors.

Our few diverging estimated parameters highlight a distinction between behavior expected from laboratory-derived relationships and realized tick behavior in the field. Within the prior effect model, mechanistic relationships were strictly informed by laboratory studies, whereas the posterior-effect model allowed those relationships to be estimated directly from field data. Our posterior estimates for nymphal questing thresholds (*∼* 85% RH) aligned well with laboratory survival expectations, whereas the model departed from the priors for both the larval humidity threshold and minimum overwinter larval-to-nymphal development rates. For the larval humidity threshold, our posterior estimates implied that larval ticks require a roughly 95% RH for 90% of the population to quest. This increased larval vulnerability is biologically plausible given larval ticks’ greater surface-area-to-volume ratio compared to nymphs and adults (Ginsberg et al., 2017). Furthermore, field evidence corroborates this heightened sensitivity, demonstrating that larvae are significantly more vulnerable to desiccation-stress than either nymphs or adults (Brunner et al., 2023). This empirical validation directly supports the plausibility of our posterior model estimates. This discrepancy highlights the distinction between behavior implied by laboratory studies (Trout Fryxell et al., 2015; Stafford, 1994) and realized field behavior. For example, although a specific environment may be sufficient for larval survival in controlled laboratory conditions, active host-seeking in the field could reasonably demand more permissive (i.e., more humid) conditions. Notably, field studies often lump larvae and nymphs together as “immature tick stages” when analyzing humidity constraints; our results suggest these stages are physiologically distinct and should be considered separately.

Our results also revealed a second departure from laboratory expectations regarding winter dynamics, as field data consistently supported a modest but nonzero overwinter larval-to-nymphal development rate. Although standard laboratory curves extrapolate that development ceases entirely below a strict temperature threshold (Ogden et al., 2004), low-level winter development remains biologically plausible. This continuous development may occur because physiological processes briefly resume during warm periods within diurnal temperature fluctuations—a dynamic masked by daily temperature averages. Alternatively, because ticks remain beneath snowpack or leaf litter in the winter, they experience microclimates significantly warmer than ambient air temperatures (Diyes et al., 2024). Similar low-level winter development has been documented in *Ixodes ricinus*, supporting the plausibility of this phenomenon within the genus (Kjellander et al., 2023).

Although winter development had minimal influence on predicted phenological timing, producing only a small increase in spring nymphal activity rather than advancing its peak emergence, relative humidity constraints fundamentally shaped regional tick phenology by narrowing the temporal window permissive for questing. Across all sites, the inclusion of humidity uniformly delayed the larval peak, with the effect being most pronounced at BLAN and SCBI. At HARV, this delay was less significant because larval activity was already tightly constrained by temperature in the early summer. The impact of humidity on nymphal phenology was more variable and contingent upon the interaction between spring humidity and temperature. At BLAN, humidity accelerated the nymphal peak; high spring temperatures allow for rapid larval-to-nymphal development, generating an accumulation of molted nymphs that are initially prohibited from questing by low humidity. Once humidity conditions permit, these individuals all initiate questing simultaneously, shifting the nymphal peak forward. Conversely, if spring temperatures are too low, as seen at HARV, low early spring development rates limit the build-up of molted nymphs, preventing the accumulation necessary to drive an earlier peak. Furthermore, if humidity is low in the spring, as seen at SCBI, the accumulated nymphs cannot begin questing until later in the season, closer to the point where temperature alone would have predicted a peak. Ultimately, the humidity-dependent model yielded more substantial improvements at the Mid-Atlantic sites (BLAN and SCBI) than at the Northeastern site (HARV), emphasizing that tick populations in the Mid-Atlantic are primarily humidity-limited, whereas those in the Northeast remain temperature-limited.

Despite the posterior model generally having higher consistency with the data than the prior .effect and no-effect models, certain site-years exhibited fairly low correlations with posterior model predictions. These discrepancies may arise from unmodeled environmental drivers, inherent process stochasticity, or the coarse temporal resolution of the NEON sampling design, generating noisy data. For example, short-term rainfall events may alter near-surface moisture conditions in ways not captured by relative humidity alone (Weiler et al., 2017), and questing activity may vary with time of day (Schulze & Jordan, 2003) or fine-scale spatial heterogeneity in vegetation (Fischhoff et al., 2019; Linske & Williams, 2024). In addition, larval counts are subject to chance dragging near “larval bombs,” in which large numbers of larvae emerge from localized egg masses, producing extremely high counts (Brunner & Ostfeld, 2008). The stochasticity of tick drags, particularly larval drags, was supported by our estimated Dirichlet-multinomial concentration parameters, suggesting moderate aggregation for nymphs and high aggregation for larvae. Despite the inherent noise in drag data, the systematic improvement in model performance over the temperature-only alternative confirms that humidity is a key component of tick phenology.

Interpretation of synchrony results requires particular care, because empirical synchrony estimates depend on activity distributions fitted to coarsely sampled NEON data. Across site-years, models incorporating the hypothesized mechanisms generally improved synchrony predictions, with clear improvement at two of the three sites. At the third site, HARV, the baseline model already produced accurate synchrony estimates in most years, so although posterior predictions in those years remained accurate, the posterior model did not improve predictions. In the remaining year, posterior estimates diverged more strongly; however, the empirical synchrony curve for that case was highly sensitive to sparse sampling, making the apparent discrepancy less informative than the broader multi-year pattern. Accordingly, trends consistent across sites and years provide a more reliable measure of model performance than outcomes from any single site-year.

In conclusion, our results identify humidity-dependent questing as a primary driver of *Ixodes scapularis* phenology and show that explicitly representing humidity as a constraint on questing is necessary to reproduce observed phenology. Unlike previous approaches that rely on fixed seasonal forcing or empirically calibrated cumulative functions, our model links phenology directly to physiological responses to climate. This mechanistic structure allows for biological interpretation and predictions beyond observed conditions, which is essential for anticipating responses under shifting climates. Accurately predicting seasonal timing is necessary to accurately predict risk. The interval between nymphal and larval activity directly shapes transmission opportunities, because it determines whether larvae encounter hosts infected by earlier-feeding nymphs and whether co-feeding transmission can occur (Randolph et al., 1996; Voordouw, 2015). These timing effects are especially important for rapidly cleared *Borrelia burgdorferi* strains, where small phenological shifts can decide pathogen persistence (States et al., 2017). Overall, our model provides a practical tool for examining how climate variation alters transmission risk and, through its improved representation of larval–nymphal synchrony, for investigating the climatic drivers of coinfection.

## Acknowledgments

This work was supported by the NIH National Institute of General Medical Sciences R35 MIRA program (grant no. R35GM143029)

## References

Arsnoe, I., Tsao, J. I., and Hickling, G. J. (2019). Nymphal Ixodes scapularis questing behavior explains geographic variation in Lyme borreliosis risk in the eastern United States. Ticks and Tick-borne Diseases, 10(3):553–563.

Belozerov, V. N., Fourie, L. J., and Kok, D. J. (2002). Photoperiodic control of developmental diapause in nymphs of prostriate ixodid ticks (Acari: Ixodidae). Experimental and Applied Acarology, 28:163–168.

Berger, K. A., Ginsberg, H. S., Dugas, K. D., Hamel, L. H., and Mather, T. N. (2014). Adverse moisture events predict seasonal abundance of Lyme disease vector ticks (Ixodes scapularis). Parasites & Vectors, 7(1):1.

Branagan, L. D. (1973). The development periods of the Ixodid tick Rhipicephalus appendiculatus Neum. under laboratory conditions. Bulletin of Entomological Research, 63(1):155–168.

Brunner, J. L., LaDeau, S., Killilea, M., Valentine, E., Schierer, M., and Ostfeld, R. S. (2023). Off-Host Survival of Blacklegged Ticks in Eastern North America: A Multistage, Multiyear, Multisite Study. Ecological Monographs, 93(3):e1572.

Brunner, J. L. and Ostfeld, R. S. (2008). Multiple causes of variable tick burdens on small-mammal hosts. Ecology, 89(8):2259–2272.

Chilton, N. B. and Bull, C. M. (1994). Influence of environmental factors on oviposition and egg development in Amblyomma limbatum and Aponomma hydrosauri (Acari: Ixodidae). International Journal for Parasitology, 24(1):83–90.

Diuk-Wasser, M. A., Vannier, E., and Krause, P. J. (2016). Coinfection by Ixodes tick-borne pathogens: Ecological, epidemiological, and clinical consequences. Trends in Parasitology, 32(1):30–42.

Diyes, C. P., Yunik, M. E. M., Dergousoff, S. J., and Chilton, N. B. (2024). Effect of snow cover on the off-host survival of Dermacentor variabilis (Acari: Ixodidae) larvae. Journal of Medical Entomology, 61(1):46–54.

Donahue, J. G., Piesman, J., and Spielman, A. (1987). Reservoir competence of white-footed mice for Lyme disease spirochetes. The American Journal of Tropical Medicine and Hygiene, 36(1):92–96.

Duffy, D. C. and Campbell, S. R. (1994). Ambient air temperature as a predictor of activity of adult Ixodes scapularis (Acari: Ixodidae). Journal of Medical Entomology, 31(1):178–180.

Eisen, L. and Eisen, R. J. (2023). Changes in the geographic distribution of the blacklegged tick, Ixodes scapularis, in the United States. Ticks and Tick-borne Diseases, 14(6):102233.

Fischhoff, I., Keesing, F., Pendleton, J., DePietro, D., Teator, M., Duerr, S. T. K., Mowry, S., Pfister, A., LaDeau, S. L., and Ostfeld, R. S. (2019). Assessing effectiveness of recommended residential yard management measures against ticks. Journal of Medical Entomology, 56(5):1420– 1427.

FitzJohn, R. and Hinsley, W. (2025). odin2: Next generation odin. R package version 0.3.19.

Gatewood, A. G., Liebman, K. A., Vourc’h, G., Bunikis, J., Hamer, S. A., Cortinas, R., Melton, F., Cislo, P., Kitron, U., Tsao, J., Barbour, A. G., Fish, D., and Diuk-Wasser, M. A. (2009). Climate and tick seasonality are predictors of Borrelia burgdorferi genotype distribution. Applied and Environmental Microbiology, 75(8):2476–2483.

Gilbert, L., Aungier, J., and Tomkins, J. L. (2014). Climate of origin affects tick (Ixodes ricinus) host-seeking behavior in response to temperature: implications for resilience to climate change? Ecology and Evolution, 4(7):1186–1198.

Ginsberg, H. S., Albert, M., Acevedo, L., Dyer, M. C., Arsnoe, I. M., Tsao, J. I., et al. (2017). Environmental factors affecting survival of immature Ixodes scapularis and implications for geographical distribution of Lyme disease: The climate/behavior hypothesis. PLoS ONE, 12(1):e0168723.

Goethert, H. K., Mather, T. N., O’Callahan, A., and Telford, III, S. R. (2023). Host-utilization differences between larval and nymphal deer ticks in northeastern U.S. sites enzootic for Borrelia burgdorferi sensu stricto. Ticks and Tick-borne Diseases, 14(6):102230.

Gray, J. S., Kahl, O., Lane, R. S., Levin, M. L., and Tsao, J. I. (2016). Diapause in ticks of the medically important Ixodes ricinus species complex. Ticks and Tick-borne Diseases, 7(5):992– 1003.

Hartig, F., Minunno, F., and Paul, S. (2023). BayesianTools: General-Purpose MCMC and SMC samplers and tools for Bayesian statistics. R package version 0.1.8.

Haven, J., Magori, K., and Park, A. W. (2012). Ecological and in host factors promoting distinct parasite life-history strategies in Lyme borreliosis. Epidemics, 4(3):152–157.

Husar, K., Pittman, D. C., Rajala, J., Mostafa, F., and Allen, L. J. S. (2024). Lyme disease models of tick-mouse dynamics with seasonal variation in births, deaths, and tick feeding. Bulletin of Mathematical Biology, 86(3):25.

Kelly, V. (2020). Cary Institute of Ecosystem Studies, Environmental Monitoring Program Data. Data set. Cary Institute.

Kjellander, P., Bergvall, U. A., Chirico, J., Ullman, K., Christensson, M., and Lindgren, P. E. (2023). Winter activity of Ixodes ricinus in Sweden. Parasites & Vectors, 16(1):229.

Kurtenbach, K., Hanincová, K., Tsao, J. I., Margos, G., Fish, D., and Ogden, N. H. (2006). Fundamental processes in the evolutionary ecology of Lyme borreliosis. Nature Reviews Microbiology, 4(9):660–669.

Levi, T., Keesing, F., Oggenfuss, K., and Ostfeld, R. S. (2015). Accelerated phenology of blacklegged ticks under climate warming. Philosophical Transactions of the Royal Society B: Biological Sciences, 370(1665):20130556.

Levin, M. L. and Fish, D. (1998). Density-dependent factors regulating feeding success of Ixodes scapularis larvae (Acari: Ixodidae). Journal of Parasitology, 84(1):36–43.

Lindsay, L. R., Barker, I. K., Surgeoner, G. A., McEwen, S. A., Gillespie, T. J., and Addison, E. M. (1998). Survival and development of the different life stages of Ixodes scapularis (Acari: Ixodidae) held within four habitats on Long Point, Ontario, Canada. Journal of Medical Entomology, 35(3):189–199.

Linske, M. A., Stafford, K. C., III, Williams, S. C., Lubelczyk, C. B., Welch, M., and Henderson, E. F. (2019). Impacts of deciduous leaf litter and snow presence on nymphal Ixodes scapularis (Acari: Ixodidae) overwintering survival in coastal New England, USA. Insects, 10(8):227.

Linske, M. A. and Williams, S. C. (2024). Evaluation of landscaping and vegetation management to suppress host-seeking Ixodes scapularis (Ixodida: Ixodidae) nymphs on residential properties in Connecticut, USA. Environmental Entomology, 53(2):268–276.

Lou, Y., Wu, J., and Wu, X. (2014). Impact of biodiversity and seasonality on Lyme-pathogen transmission. Theoretical Biology and Medical Modelling, 11:50.

Ludwig, A., Ginsberg, H. S., Hickling, G. J., and Ogden, N. H. (2016). A dynamic population model to investigate effects of climate and climate-independent factors on the lifecycle of Amblyomma americanum (Acari: Ixodidae). Journal of Medical Entomology, 53(1):99–115.

MacDonald, H., Akçay, E., and Brisson, D. (2021). The role of host phenology for parasite transmission. Theoretical Ecology, 14(1):123–143.

Mays, S. E., Houston, A. E., and Trout Fryxell, R. T. (2016). Comparison of novel and conventional methods of trapping ixodid ticks in the southeastern U.S.A. Medical and Veterinary Entomology, 30(2):123–134.

Mount, G. A., Haile, D. G., and Daniels, E. (1997). Simulation of blacklegged tick (Acari: Ixodidae) population dynamics and transmission of Borrelia burgdorferi. Journal of Medical Entomology, 34(4):461–484.

Musa, S. S. and Gumel, A. B. (2026). Could global warming cause a range expansion or shift of Lyme disease in the U.S. state of Maryland? A mathematical modelling approach. Royal Society Open Science, 13:251672.

National Ecological Observatory Network (NEON) (2024a). Relative humidity (DP1.00098.001), provisional data. Retrieved November 10, 2024.

National Ecological Observatory Network (NEON) (2024b). Triple aspirated air temperature (DP1.00003.001), provisional data. Retrieved June 10, 2024.

National Ecological Observatory Network (NEON) (2025). Ticks sampled using drag cloths (DP1.10093.001), RELEASE-2022.

National Institute of Allergy and Infectious Diseases (2018). Lyme disease coinfection.

Ogden, N. H., Ben Beard, C., Ginsberg, H. S., and Tsao, J. I. (2021). Possible effects of climate change on ixodid ticks and the pathogens they transmit: Predictions and observations. Journal of Medical Entomology, 58(4):1536–1545.

Ogden, N. H., Bigras-Poulin, M., O’Callaghan, C. J., Barker, I. K., Lindsay, L. R., Maarouf, A., Smoyer-Tomic, K. E., Waltner-Toews, D., and Charron, D. (2005). A dynamic population model to investigate effects of climate on geographic range and seasonality of the tick Ixodes scapularis. International Journal for Parasitology, 35(4):375–389.

Ogden, N. H., Lindsay, L. R., Charron, D., Beauchamp, G., Maarouf, A., O’Callaghan, C. J., Waltner-Toews, D., and Barker, I. K. (2004). Investigation of the relationships between temperature and development rates of the tick Ixodes scapularis (Acari: Ixodidae) in the laboratory and field. Journal of Medical Entomology, 41(4):622–633.

Ogden, N. H., Lindsay, L. R., and Leighton, P. A. (2013). Predicting the rate of invasion of the agent of Lyme disease Borrelia burgdorferi. Journal of Applied Ecology, 50:510–518.

Ostfeld, R. S. and Brunner, J. L. (2015). Climate change and Ixodes tick-borne diseases of humans. Philosophical Transactions of the Royal Society B: Biological Sciences, 370(1665):20140051.

Peavey, C. A. and Lane, R. S. (1996). Field and laboratory studies on the timing of oviposition and hatching of the western black-legged tick, Ixodes pacificus (Acari: Ixodidae). Experimental and Applied Acarology, 20(12):695–711.

Perret, J. L., Guerin, P. M., Diehl, P. A., Vlimant, M., and Gern, L. (2003). Darkness induces mobility, and saturation deficit limits questing duration, in the tick Ixodes ricinus. Journal of Experimental Biology, 206(11):1809–1815.

Qviller, L., Grøva, L., Viljugrein, H., Klingen, I., and Mysterud, A. (2014). Temporal pattern of questing tick Ixodes ricinus density at differing elevations in the coastal region of western Norway. Parasites & Vectors, 7:179.

Randolph, S. E. (1997). Abiotic and biotic determinants of the seasonal dynamics of the tick Rhipicephalus appendiculatus in South Africa. Medical and Veterinary Entomology, 11(1):25– 37.

Randolph, S. E., Gern, L., and Nuttall, P. A. (1996). Co-feeding ticks: Epidemiological significance for tick-borne pathogen transmission. Parasitology Today, 12(12):472–479.

Randolph, S. E., Green, R. M., Hoodless, N. P., and Peacey, M. F. (2002). An empirical quantitative framework for the seasonal population dynamics of the tick Ixodes ricinus. International Journal for Parasitology, 32(8):979–989.

Rodgers, S. E., Zolnik, C. P., and Mather, T. N. (2007). Duration of exposure to suboptimal atmospheric moisture affects nymphal blacklegged tick survival. Journal of Medical Entomology, 44(2):372–375.

Sakamoto, J. M., Goddard, J., and Rasgon, J. L. (2014). Population and demographic structure of Ixodes scapularis Say in the eastern United States. PLoS ONE, 9(7):e101389.

Sambado, S., MacDonald, A. J., Swei, A., and Briggs, C. J. (2024). Climate-associated variation in the within-season dynamics of juvenile ticks in California. Ecosphere, 15(11):e70064.

Schauber, E. M. and Ostfeld, R. S. (2002). Modeling the Effects of Reservoir Competence Decay and Demographic Turnover in Lyme Disease Ecology. Ecological Applications, 12(4):1142– 1162.

Schulze, T. L. and Jordan, R. A. (2003). Meteorologically mediated diurnal questing of Ixodes scapularis and Amblyomma americanum (Acari: Ixodidae) nymphs. Journal of Medical Entomology, 40(4):395–402.

Stafford, III, K. C. (1994). Survival of immature Ixodes scapularis (Acari: Ixodidae) at different relative humidities. Journal of Medical Entomology, 31(2):310–314.

States, S. L., Huang, C. I., Davis, S., Tufts, D. M., and Diuk-Wasser, M. A. (2017). Co-feeding transmission facilitates strain coexistence in Borrelia burgdorferi, the Lyme disease agent. Epidemics, 19:33–42.

Steer, N. C., Ramsay, P. M., and Franco, M. (2019). nlstimedist: An R package for the biologically meaningful quantification of unimodal phenology distributions. Methods in Ecology and Evolution, 10(11):1934–1940.

T. Fryxell, R. T., Moore, J. E., Collins, M. D., Kwon, Y., Jean-Philippe, S. R., Schaeffer, S. M., Odoi, A., Kennedy, M., and Houston, A. E. (2015). Habitat and vegetation variables are not enough when predicting tick populations in the southeastern United States. PLoS ONE, 10(12):e0144092.

Tagliapietra, V., Rosà, R., Arnoldi, D., Cagnacci, F., Capelli, G., Montarsi, F., Hauffe, H. C., and Rizzoli, A. (2011). Saturation deficit and deer density affect questing activity and local abundance of Ixodes ricinus (Acari, Ixodidae) in Italy. Veterinary Parasitology, 183(1-2):114–124.

Thomas, V., Anguita, J., Barthold, S. W., and Fikrig, E. (2001). Coinfection with Borrelia burgdorferi and the agent of human granulocytic ehrlichiosis alters murine immune responses, pathogen burden, and severity of Lyme arthritis. Infection and Immunity, 69(5):3359–3371.

Vail, S. C., Smith, G. J., and Lord, C. C. (1994). Population biology of Ixodes scapularis, the vector of Lyme disease in the Eastern and North Central United States. In Scott, M. E. and Smith, G. C., editors, Parasitic and Infectious Diseases: Epidemiology and Ecology, pages 263–278. Academic Press.

Vail, S. G. and Smith, G. (2002). Vertical movement and posture of blacklegged tick (Acari: Ixodidae) nymphs as a function of temperature and relative humidity in laboratory experiments. Journal of Medical Entomology, 39(6):842–846.

Venables, W. N. and Ripley, B. D. (2002). Modern applied statistics with S. Springer, 4th edition.

Volk, M. R., Lubelczyk, C. B., Johnston, J. C., Levesque, D. L., and Gardner, A. M. (2022). Microclimate conditions alter Ixodes scapularis (Acari: Ixodidae) overwinter survival across climate gradients in Maine, United States. Ticks and Tick-borne Diseases, 13(1):101872.

Voordouw, M. J. (2015). Co-feeding transmission in Lyme disease pathogens. Parasitology, 142(2):290–302.

Wallace, D., Ratti, V., Kodali, A., Winter, J. M., Ayres, M. P., Chipman, J. W., Aoki, C. F., Osterberg, E. C., Silvanic, C., Partridge, T. F., and Webb, M. J. (2019). Effect of rising temperature on Lyme disease: Ixodes scapularis population dynamics and Borrelia burgdorferi transmission and prevalence. Canadian Journal of Infectious Diseases and Medical Microbiology, 2019:9817930.

Weiler, M., Duscher, G. G., Wetscher, M., and Walochnik, J. (2017). Tick abundance: A one year study on the impact of flood events along the banks of the river Danube, Austria. Experimental and Applied Acarology, 71(2):151–157.

Winter, J. M., Partridge, T. F., Wallace, D., Chipman, J. W., Ayres, M. P., Osterberg, E. C., and Dekker, E. R. (2021). Modeling the sensitivity of blacklegged ticks (Ixodes scapularis) to temperature and land cover in the northeastern United States. Journal of Medical Entomology, 58(1):416–427.

Wolf, M. J., Watkins, H. R., and Schwan, W. R. (2020). Ixodes scapularis: Vector to an increasing diversity of human pathogens in the upper Midwest. WMJ: Official Publication of the State Medical Society of Wisconsin, 119(1):16–21.

Wood, S. N. (2011). Fast stable restricted maximum likelihood and marginal likelihood estimation of semiparametric generalized linear models. Journal of the Royal Statistical Society: Series B (Statistical Methodology), 73(1):3–36.

Wu, X., Duvvuri, V. R., Lou, Y., Ogden, N. H., Pelcat, Y., and Wu, J. (2013). Developing a temperature-driven map of the basic reproductive number of the emerging tick vector of Lyme disease Ixodes scapularis in Canada. Journal of Theoretical Biology, 319:50–61.

Wu, X., Duvvuri, V. R. S. K., and Wu, J. (2010). Modeling dynamical temperature influence on tick Ixodes scapularis population. In International Congress on Environmental Modelling and Software, Ottawa, Canada.

Yuval, B. and Spielman, A. (1990). Duration and regulation of the developmental cycle of Ixodes dammini (Acari: Ixodidae). Journal of Medical Entomology, 27(2):196–201.

