## Supplemental Material for "Refining mechanistic models to better predict larval and nymphal activity patterns of *Ixodes scapularis*"

### 1 Introduction

#### 1.1 Limitations of accumulating daily development rates

A common strategy in tick life cycle models is to estimate development duration by accumulating daily temperature-dependent rates until a cumulative threshold of one is reached, then the effective development rate is defined as the reciprocal of the estimated duration (Wu et al., 2013; Lou et al., 2014). Although this formulation reproduces observed nymphal activity under contemporary temperate climates in North America, it introduces structural inconsistencies that remove the biological interpretation and predictive utility of the model. Specifically, the approach loses mechanistic interpretability, real-time forecasting, and out-of-sample predictive capacity and masks problems with the temperature-only formulation.

*Loss of mechanistic interpretation*
The accumulation technique disconnects modeled development from the underlying physiology. Since development is calculated only after the full stage duration is known, individual days are assigned developmental rates based on future environmental conditions rather than the day's thermal suitability. As a result, biologically impossible conditions can still contribute to modeled progression. For example, consider a hypothetical period beginning with a day well below the developmental threshold, followed by ten days at the thermal optimum (ie., a development rate of 1/10 each day). Under the accumulation framework, development would be complete after eleven total days. Using the accumulation technique, the initial day is then assigned a development rate of 1/11, implying that nearly optimal development (characterized by a daily development rate of 1/10) can occur on a day with temperature far below the development threshold.

*Loss of real-time forecasting*
As the developmental rate using this technique depends on the eventual stage duration, the model cannot determine progression at a given time point with-

out knowledge of future environmental conditions. Development on day $t$ is contingent on temperatures occurring after day $t$, introducing an inherent dependence on future climate trajectories. This structure prevents the model from functioning as a true forecasting framework, limiting the model's utility for operational public health applications such as early-warning systems, seasonal risk forecasting, or forward climate projections.

*Loss of out-of-sample predictive capacity*
The accumulation framework achieves temporal agreement with observed phenology by implicitly relying on historically stable seasonal temperature structure. Specifically, accurate developmental timing relies on a typical climate pattern of a cold winter followed by a spring warm up, allowing the winter to "borrow" development from the spring. As a consequence, the model's predictive performance is dependent on historical climate patterns. When applied to a truly novel environment, for example, one that does not conform to the warm following cold pattern, the model will likely no longer generate accurate predictions. The model therefore lacks robust out-of-sample predictive capacity.

*Masking deficiencies in temperature-only ODE formulations*
A standard temperature-driven ODE formulation without retrospective accumulation predicts nymphal emergence substantially later than observed empirically (Winter et al., 2020; Wu et al., 2013). The accumulation framework compensates for this mismatch by effectively borrowing spring development, artificially accelerating nymphal emergence timing from the mid summer to mid spring. Critically, this apparent improvement does not arise because the underlying biology has been correctly specified. Instead, the retrospective method introduces a mathematical artifact that shifts activity peaks forward in time, creating the appearance of accurate phenology. In practice, the method produces a correct fit to observed data but for the wrong mechanistic reasons, therefore masking the absence of necessary additional biological drivers.

### 2 Materials and methods

#### 2.1 Original model description

The model consists of 12 discrete tick life stages. Ticks begin as eggs ($E$). Eggs either develop into hardening larvae ($L_H$) at a temperature-dependent development rate of $\rho_{el}(T(t))$ (where $T(t)$ is the temperature on day $t$) or die at a rate $\mu_e$. Hardening larvae represent individuals that are immediately post-molt and therefore are unable to feed. Hardening larvae develop into larvae that are ready to feed at a constant rate of $\rho_h$ or die at a constant rate $\mu_{hl}$. Larvae that are ready to feed ($L_R$) then transition to feeding larvae once temperature conditions allow for questing and a host is available, governed by the probability of questing rate, $\theta^l(T(t))$, and host-finding rate, $\lambda_{rl}(H_i)$, or die at rate $\mu_{rl}$. Feeding larvae ($L_F$) next develop into engorged larvae at a constant rate $\rho_{fl}$ or die at a density-dependent mortality rate $\mu_{fl}(L_F)$. Engorged larvae ($L_E$)

develop into nymphs that are ready to feed at a temperature dependent rate $\rho_{ln}(T(t))$ or die at rate $\mu_{el}$. Similar to larvae that are ready to feed, nymphs that are ready to feed ($N_R$) transition to feeding nymphs based on host-finding probability $\lambda_{rn}(H_i)$ and questing rate $\theta^n(T(t))$ or die at rate $\mu_{rn}$. Fed nymphs become engorged ($N_E$) at constant rate $\rho_{fn}$ or die at density-dependent rate $\mu_{fn}(N_F)$. Nymphs transition to adults that are ready to feed ($A_R$) in the same manner as described for larvae, with a development rate $\rho_{na}(T(t))$ or die at a rate of $\mu_{en}$. Since only female adults take a blood meal, one half of adults that are ready to feed become feeding adults ($A_F$) based on the host-finding rate $\lambda_{ra}(H_A)$ and questing rate $\theta^a(T(t))$. Feeding adults die at a density-dependent mortality rate, $\mu_{fa}(A_F)$, or transition to engorged adults ($A_E$) at a constant rate $\rho_{fa}$. Engorged adults either develop into gravid adults ($A_G$) at a temperature-dependent rate $\rho_{ae}(T(t))$ or die at rate $\mu_{ea}$. Finally, gravid adults lay a density-dependent number of eggs, $f(A_F)\varepsilon$, at a rate $\rho_O$. All development rates are defined as in Ogden 2005 with life-stage specific parameters provided in Supplemental Table 1. Density-dependent mortality was modeled as in Ogden 2005 where

$$\mu_{fs} = a + \left(0.049 \log\left(1.01 + \frac{L_s}{H}\right)\right) \tag{1}$$

for each life stage $s$ (Stage specific parameters provided in Table S2). Similarly, questing functions $\theta(T(t))$ were modeled as in Ogden 2005 such that,

$$\theta_l(T(t)) = \begin{cases} 0 & \text{if } T(t) \leq 0 \\ \max(0, -0.291 + 0.002T(t)^2) & \text{if } 0 < T(t) < 26.6 \\ \max(0, 3.111 - 0.084T(t)) & \text{if } T(t) \geq 26.6 \end{cases} \tag{2}$$

$$\theta_n(T(t)) = \max(0, -0.060 + 0.114T(t) - 0.003T(t)^2) \tag{3}$$

$$\theta_a(T(t)) = \max(0, -0.637 + 0.379T(t) - 0.021T(t)^2) \tag{4}$$

Host finding rates were also modeled as in Ogden 2005 with

$$\lambda_{rs}(H_s) = aH_s^b, \tag{5}$$

with parameters for each life stage $s$ provided in Table S3. Photoperiod-driven diapause was incorporated as in the original model by specifying that $\rho(T(t)) = 0$ when $t > 180$, representing the cessation of development as day length shortens beyond the summer solstice. The full system of equations for the original model is provided:

$$
\begin{aligned}
\frac{dA_G}{dt} &= \rho_{ae}(T(t))A_E - \rho_o A_G \\
\frac{dE}{dt} &= A_G f(A_F)\varepsilon - (\rho_{el}(T(t)) + \mu_e)E \\
\frac{dL_H}{dt} &= \rho_{el}(T(t))E - (\rho_h + \mu_{hl})L_H \\
\frac{dL_R}{dt} &= \rho_h L_H - (\lambda_{rl}(H_i)\theta^l(T(t)) + \mu_{rl})L_R \\
\frac{dL_F}{dt} &= \lambda_{rl}(H_i)\theta^l(T(t))L_R - (\rho_{fl} + \mu_{fl}(L_F))L_F \\
\frac{dL_E}{dt} &= \rho_{fl}L_F - (\mu_{el} + \rho_{ln}(T(t)))L_E \\
\frac{dN_R}{dt} &= \rho_{ln}(T(t))L_E - (\lambda_{rn}(H_i)\theta^n(T(t)) + \mu_{rn})N_R \\
\frac{dN_F}{dt} &= \lambda_{rn}(H_i)\theta^n(T(t))N_R - (\rho_{fn} + \mu_{fn}(N_F))N_F \\
\frac{dN_E}{dt} &= \rho_{fn}N_F - \rho_{na}(T(t))N_E - \mu_{en}N_E \\
\frac{dA_R}{dt} &= \rho_{na}(T(t))N_E - (\lambda_{ra}(H_A)\theta^a(T(t)) + \mu_{ra})A_R \\
\frac{dA_F}{dt} &= \frac{1}{2}\lambda_{ra}(H_A)\theta^a(T(t))A_R - (\rho_{fa} + \mu_{fa}(A_F))A_F \\
\frac{dA_E}{dt} &= \rho_{fa}A_F - (\rho_{ae}(T(t)) + \mu_{ea})A_E
\end{aligned}
$$

Table 5: Description of priors and justifications

| **Parameter** | **Description** | **Prior** | **Justification** |
|---|---|---|---|
| $\overline{\delta}$ | Reparameterized shared between site mean for minimum larval-to-nymphal overwinter development rate across sites | LogNormal(0.8, 0.7) | Strictly positive, strongly concentrated around 0 reflecting prior expectation of no effect |
| $\Phi_\delta$ | Site level variance around the minimum larval-to-nymphal overwinter development rate | Exp(1/0.7) | Strictly positive, shrinkage towards mean |

Continued on next page...

Table 5: Description of priors and justifications (CONTINUED)

| Parameter | Description | Prior | Justification |
|---|---|---|---|
| $h_n$ | Shared between site mean for Nymphal threshold | Normal(30.5, 2) | 95% CI indicates that 90% of nymphal ticks will be active between 72% and 92% RH with median 82% |
| $\Phi_{hn}$ | Site level variance around the nymphal threshold | Exp(1/0.5) | Strictly positive, shrinkage towards mean |
| $h_l/h_a$ | Reparameterized shared between site mean of larval/adult threshold | LogNormal(0.1, 0.5) | Strictly positive, most of mass around 1 reflecting modest difference between lifestage humidity thresholds |
| $\Phi_{hl}/\Phi_{ha}$ | Site level variance around reparameterized larval/adult thresholds | Exp(1/0.5) | Strictly positive, shrinkage towards mean |
| $\delta_\eta$ | Site level minimum larval-to-nymphal overwinter development | LogNormal($\log(\bar{\delta})$, $\log(\Phi_\delta)$) | Between site pooling |
| $h_{n\eta}$ | Site level nymphal humidity threshold | Normal($\bar{h}$, $\Phi_{hn}$) | Between site pooling |
| $h_{l\eta}/h_{a\eta}$ | Reparameterized site level larval/adult threshold | LogNormal($\log(\bar{h}_\eta)$, $\log(\Phi_\eta)$) | Between site pooling |
| $\alpha_{sy\eta}$ | Dirichlet-Multinomial concentration parameter | LogNormal(2, 1) | Expectation of overdispersion in tick count data |

### 3 Results

Table 1: Original model parameters: Development, feeding, and reproduction

| **Parameter** | **Definition** | **Value** | **Source** |
|---|---|---|---|
| $\rho_o$ | Oviposition rate | 1/1 | Vail et al., 1994 |
| $\rho_h$ | Hardening rate | 1/21 | Vail et al., 1994 |
| $\rho_{fl}$ | Larvae feeding rate | 1/3 | Vail et al., 1994 |
| $\rho_{fn}$ | Nymph feeding rate | 1/5 | Vail et al., 1994 |
| $\rho_{fa}$ | Adult feeding rate | 1/10 | Vail et al., 1994 |
| $\rho_{ae}(T(t))$ | Pre-oviposition rate | a = 1300; b = -1.42 | Ogden et al., 2004 |
| $\rho_{el}(T(t))$ | Pre-eclosion rate | a = 34,234; b= -2.27 | Ogden et al., 2004 |
| $\rho_{ln}(T(t))$ | Larvae to nymph development rate | a = 101,181; b = -2.55 | Ogden et al., 2004 |
| $\rho_{na}(T(t))$ | Nymph to adult development rate | a = 1596; b = -1.21 | Ogden et al., 2004 |
| $f(A_F)$ | Density-dependent reduction in fecundity of egg-laying adults | a = 0.01; b = 0.04; c= 1.01 | Levin and Fish (1998) |
| $\varepsilon$ | Per capita egg production | 3000 | Mount et al., 1997 |

Table 2: Original model parameters: Mortality rates

| **Parameter** | **Definition** | **Value** | **Source** |
|---|---|---|---|
| $\mu_e$ | Mortality rate of eggs | 0.002 | Lindsay et al., 1998 |
| $\mu_{hl}$ | Mortality rate of hardening larvae | 0.006 | Lindsay et al., 1998 |
| $\mu_{rl}$ | Mortality rate of larvae ready to feed | 0.006 | Lindsay et al., 1998 |
| $\mu_{rn}$ | Mortality rate of nymphs ready to feed | 0.006 | Lindsay et al., 1998 |
| $\mu_{ra}$ | Mortality rate of adults ready to feed | 0.006 | Lindsay et al., 1998 |
| $\mu_{el}$ | Mortality rate of engorged larvae | 0.003 | Lindsay et al., 1998 |
| $\mu_{en}$ | Mortality rate of engorged nymphs | 0.002 | Lindsay et al., 1998 |
| $\mu_{ea}$ | Mortality rate of engorged adults | 0.0001 | Lindsay et al., 1998 |
| $\mu_{fl}(L_F)$ | Density-dependent mortality of feeding larvae | a = 0.65 | Levin and Fish (1998) |
| $\mu_{fn}(N_F)$ | Density-dependent mortality of feeding nymphs | a = 0.55 | Levin and Fish (1998) |
| $\mu_{fa}(A_F)$ | Density-dependent mortality of feeding adults | a = 0.5 | Levin and Fish (1998) |

Table 3: Original model parameters: Host-finding and questing

| **Parameter** | **Definition** | **Value** | **Source** |
|---|---|---|---|
| $H_i$ | Number of hosts for immature ticks | 2000 | Assumed |
| $H_A$ | Number of hosts for adult ticks | 200 | Assumed |
| $\theta^l(T(t))$ | Questing probability of larvae | a = -0.2912; b = 0.0016; c = 3.111; d = -0.084 e = -48.6; f = 0.6 | Vail and Smith (2002), Ginsberg (2017), Rodgers (2007) |
| $\theta^n(T(t))$ | Questing probability of nymphs | a = -0.0034; b = 0.1140; c = -0.0600; d = -30.5, e = 0.4 | Gilbert (2014), Ginsberg (2017), Rodgers (2007) |
| $\theta^a(T(t))$ | Questing probability of adults | a = -0.6371; b = 0.3792; c = -0.0217; d = -30.5, e = 0.4 | Duffy and Campbell (1994), Ginsberg (2017), Rodgers (2007) |
| $\lambda_{rl}$ | Daily host-finding probability for larvae | a = 0.0013; b = 0.515 | Mount et al., 1997 |
| $\lambda_{rn}$ | Daily host-finding probability for nymphs | a = 0.0013; b = 0.515 | Mount et al., 1997 |
| $\lambda_{ra}$ | Daily host-finding probability for adults | a = 0.0086; b = 0.515 | Mount et al., 1997 |

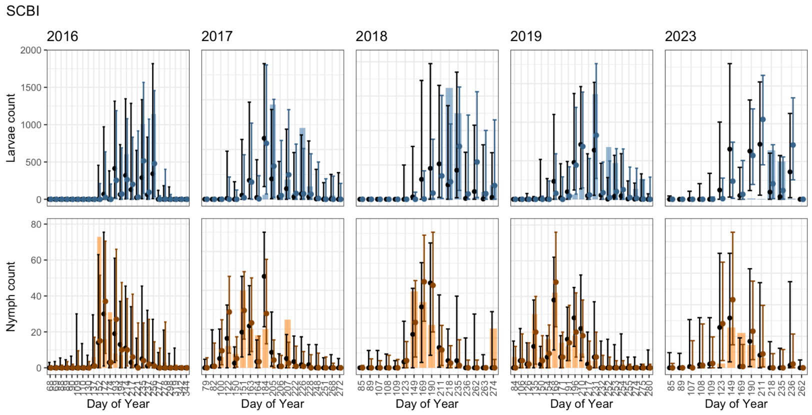


Figure 1: **Prior/Posterior predictive checks for SCBI.** Larval data and model predictions in top row, nymphal data and model predictions in bottom row. Posterior estimates and CIs in color, prior estimates and CIs in black.

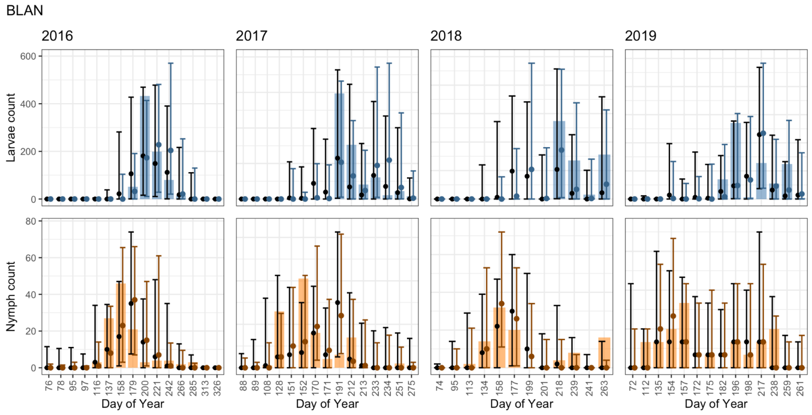


Figure 2: **Prior/Posterior predictive checks for BLAN.** Larval data and model predictions in top row, nymphal data and model predictions in bottom row. Posterior estimates and CIs in color, prior estimates and CIs in black.

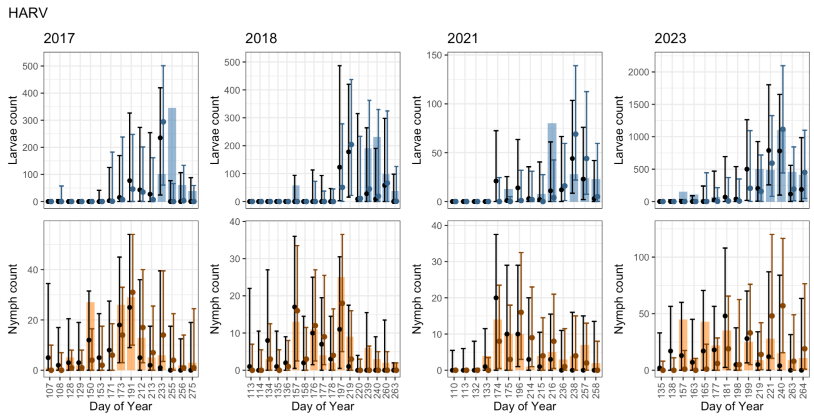


Figure 3: **Prior/Posterior predictive checks for HARV.** Larval data and model predictions in top row, nymphal data and model predictions in bottom row. Posterior estimates and CIs in color, prior estimates and CIs in black.

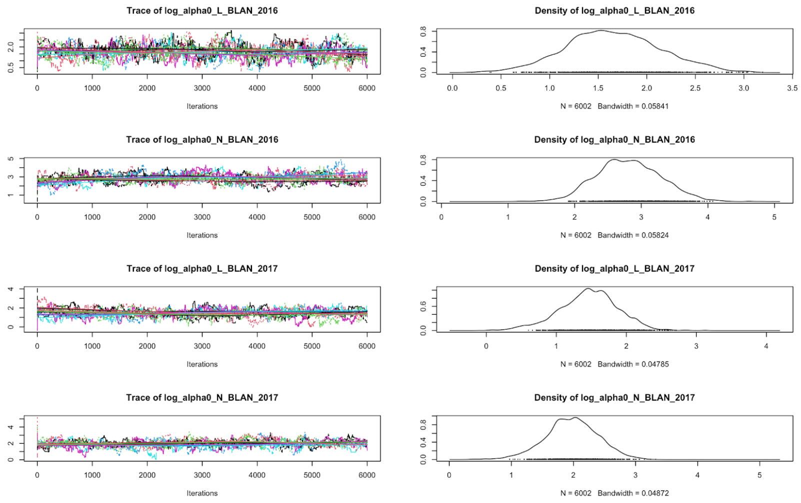


Figure 4: **Sample trace plots and density plots demonstrating chain convergence.**

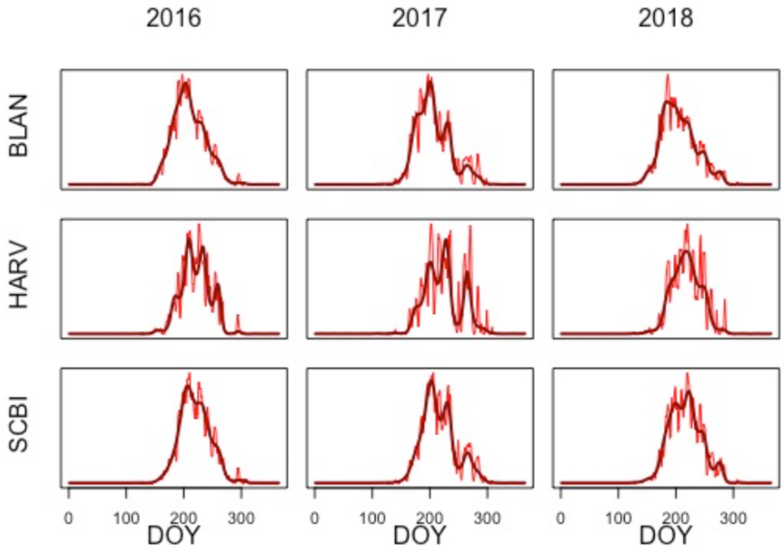


Figure 5: **Sample larval activity predictions for each site and year from the prior-effect model and fitted GAMs.** Model predictions in red and fitted GAMs used for analysis in dark red.

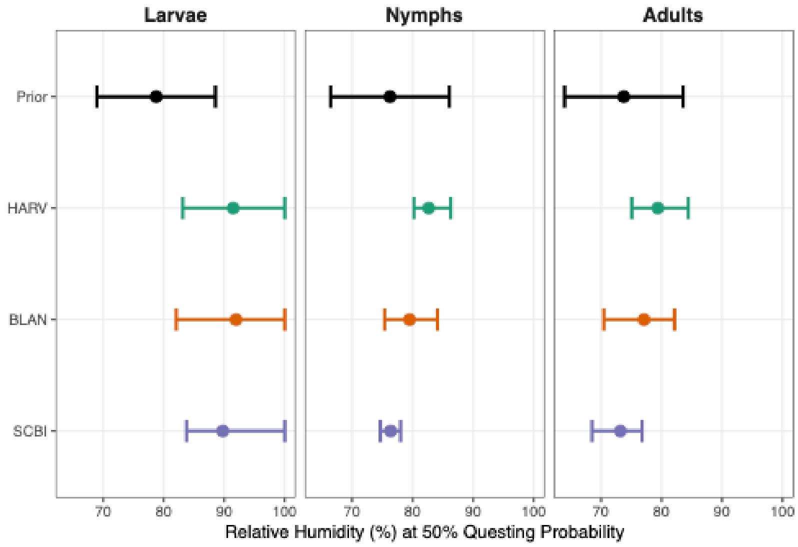


Figure 6: **Posterior estimates for effect of humidity from each site and 95% credibility intervals compared to the prior distribution.**

Table 4: Introduced model parameters in modified model

| **Parameter** | **Definition** | **Value** | **Source** |
|---|---|---|---|
| $h_a$ | Threshold for humidity on questing for adults | fitted | fitted |
| $h_n$ | Threshold for humidity on questing for nymphs | fitted | fitted |
| $h_l$ | Threshold for humidity on questing for larvae | fitted | fitted |
| $\delta$ | Minimum development rate | fitted | fitted |
| $\alpha_{ly\eta}$ | DM concentration parameter for larvae | fitted | fitted |
| $\alpha_{ny\eta}$ | DM concentration parameter for nymphs | fitted | fitted |

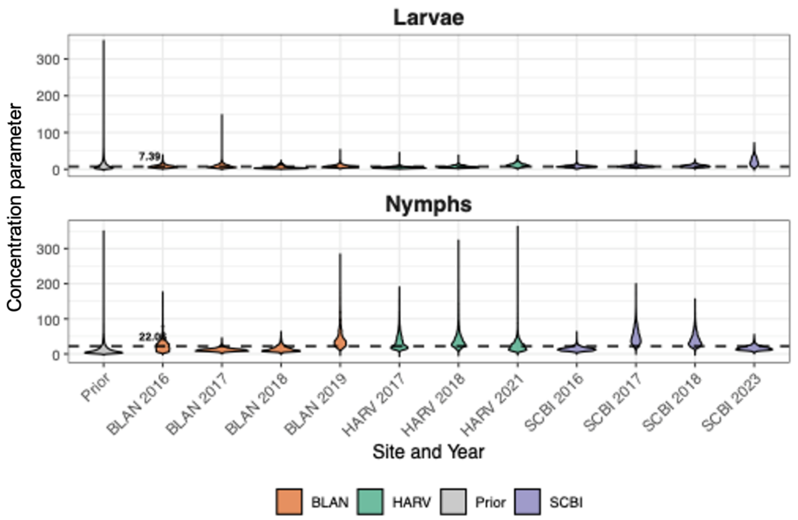


Figure 7: **Violin plots displaying full posterior distributions of concentration parameters across all specific site-years.** Labeled horizontal line displays overall mean value.

Table 6: Rank correlation for each year and life stage AT BLAN between data and the no-effect, prior-effect, and posterior-effect models

| Site | Year | Stage | No effect | prior | posterior |
|---|---|---|---|---|---|
| BLAN | 2016 | L | 0.87 | 0.92 | 0.98 |
| BLAN | 2016 | N | 0.53 | 0.87 | 0.88 |
| BLAN | 2017 | L | 0.50 | 0.66 | 0.87 |
| BLAN | 2017 | N | 0.68 | 0.67 | 0.63 |
| BLAN | 2018 | L | 0.09 | 0.46 | 0.67 |
| BLAN | 2018 | N | 0.49 | 0.55 | 0.59 |
| BLAN | 2019 | L | 0.60 | 0.81 | 0.87 |
| BLAN | 2019 | N | 0.39 | 0.49 | 0.52 |

Table 7: Rank correlation for each year and life stage AT HARV between data and the no-effect, prior-effect, and posterior-effect models

| Site | Year | Stage | No effect | prior | posterior |
|---|---|---|---|---|---|
| HARV | 2017 | L | 0.04 | 0.04 | 0.17 |
| HARV | 2017 | N | 0.26 | 0.35 | 0.60 |
| HARV | 2018 | L | 0.32 | 0.28 | 0.39 |
| HARV | 2018 | N | 0.54 | 0.50 | 0.65 |
| HARV | 2021 | L | 0.38 | 0.45 | 0.60 |
| HARV | 2021 | N | 0.58 | 0.70 | 0.63 |
| HARV | 2023 | L | 0.26 | 0.53 | 0.55 |
| HARV | 2023 | N | 0.42 | 0.54 | 0.37 |

Table 8: Variance of distribution of estimated synchrony for each site-year in the prior and posterior model and the percent reduction associated with the posterior model

| **Site** | **Year** | **Prior variance** | **Posterior variance** | **Percent reduction** |
|---|---|---|---|---|
| BLAN | 2016 | 0.0180 | 0.0130 | 27.6 |
| BLAN | 2017 | 0.0108 | 0.00958 | 11.5 |
| BLAN | 2018 | 0.00721 | 0.0120 | -66.4 |
| BLAN | 2019 | 0.0178 | 0.0115 | 35.1 |
| HARV | 2017 | 0.0209 | 0.00710 | 66.1 |
| HARV | 2018 | 0.0142 | 0.00179 | 87.3 |
| HARV | 2021 | 0.0143 | 0.00373 | 74.0 |
| SCBI | 2016 | 0.0226 | 0.00267 | 88.2 |
| SCBI | 2017 | 0.00641 | 0.000392 | 93.9 |
| SCBI | 2018 | 0.0128 | 0.000676 | 94.7 |
| SCBI | 2019 | 0.0137 | 0.00102 | 92.6 |
| SCBI | 2023 | 0.0351 | 0.00259 | 92.6 |

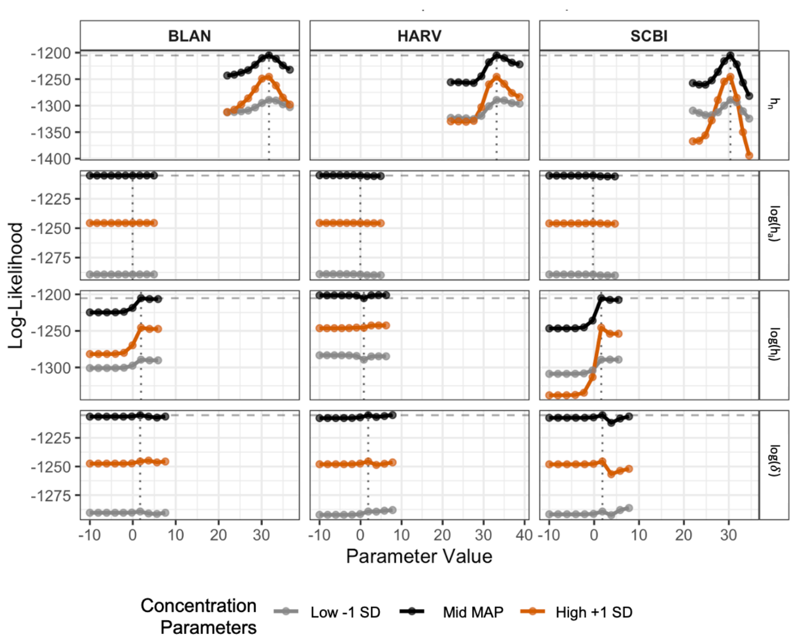


Figure 8: **Log-likelihood profiles from a one-at-a-time sensitivity analysis for mechanistic parameters.** Vertical dashed lines indicate the maximum a posteriori (MAP) estimate derived from the posterior distribution, horizontal dashed lines indicate the maximum likelihood estimate for each parameter. Alignment between the peak of a likelihood profile and its corresponding MAP estimate demonstrates that the parameter's optimal value is driven by the data, independent of the prior distribution

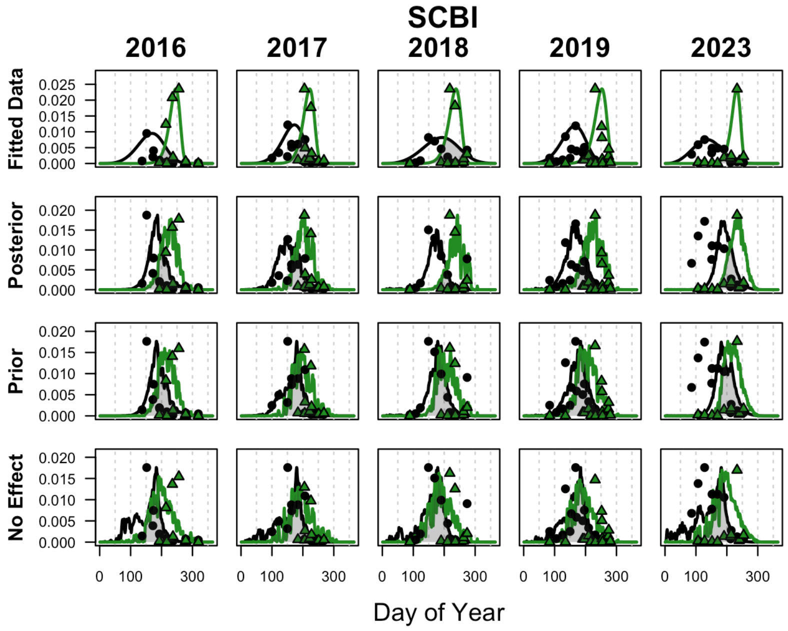


Figure 9: **Comparison of data, statistical fit, and mechanistic model predictions for SCBI.** In all panels, points represent observed field data and solid lines represent fitted activity curves for nymphs (black) and larvae (green). The top row displays empirical curves derived from statistical fits (Weibull distributions), representing the best estimate of true phenological peaks. Subsequent rows display median predictions from the posterior and prior models, alongside predictions from the no-effect model. The gray shaded regions emphasize seasonal synchrony.

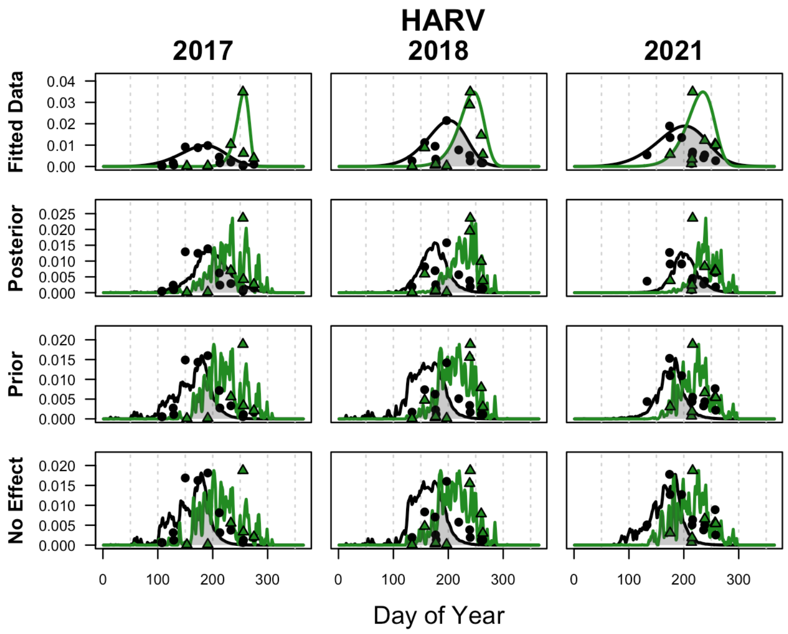


Figure 10: **Comparison of data, statistical fit, and mechanistic model predictions for HARV.** In all panels, points represent observed field data and solid lines represent fitted activity curves for nymphs (black) and larvae (green). The top row displays empirical curves derived from statistical fits (Weibull distributions), representing the best estimate of true phenological peaks. Subsequent rows display median predictions from the posterior and prior models, alongside predictions from the no-effect model. The gray shaded regions emphasize seasonal synchrony.

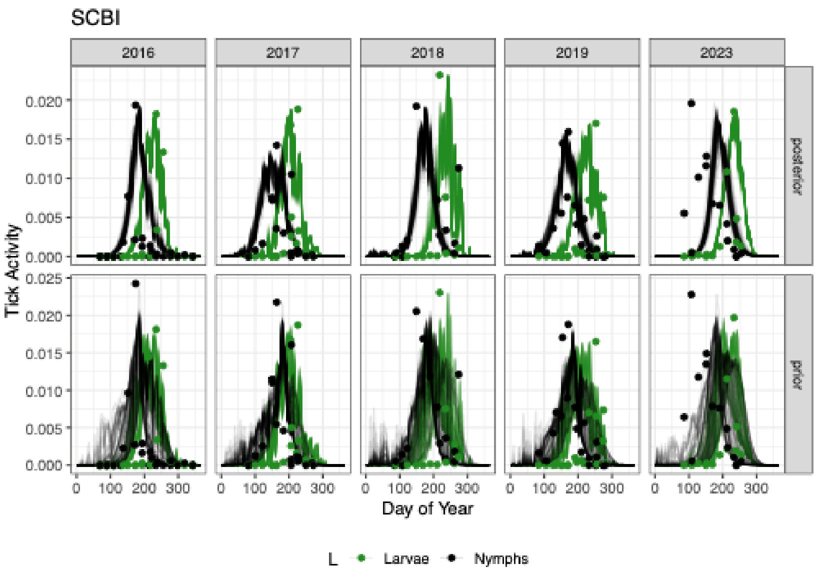


Figure 11: **Full prior and posterior predictive distribution for SCBI models.**

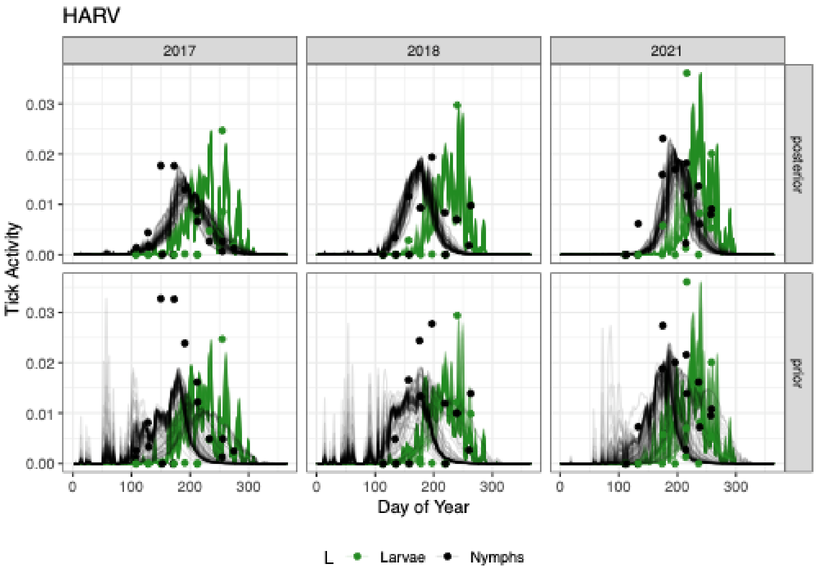


Figure 12: **Full prior and posterior predictive distribution for HARV models.**

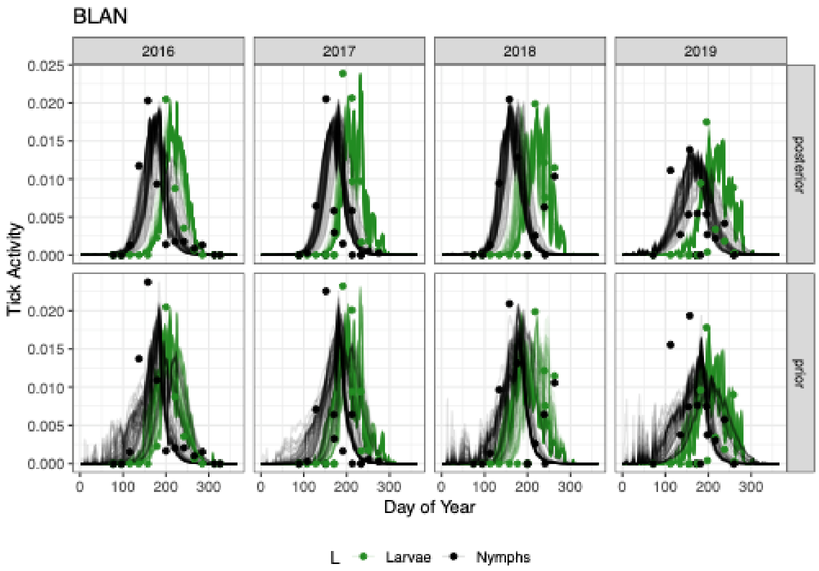


Figure 13: **Full prior and posterior predictive distribution for BLAN models.**

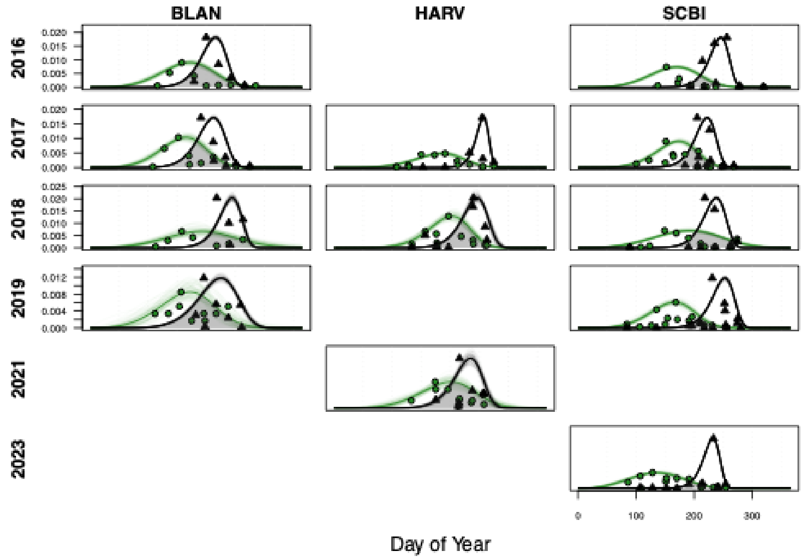


Figure 14: **Uncertainty in estimated empirical activity curves quantified by applying a Laplace approximation to the Weibull parameters.**
